# IFT-A Structure Reveals Carriages for Membrane Protein Transport into Cilia

**DOI:** 10.1101/2022.08.09.503213

**Authors:** Sophie J. Hesketh, Aakash G. Mukhopadhyay, Dai Nakamura, Katerina Toropova, Anthony J. Roberts

**Author notes:** These authors contributed equally (Sophie J. Hesketh, Aakash G. Mukhopadhyay).

## Abstract

Intraflagellar transport (IFT) trains are molecular machines that traffic proteins between cilia and the cell body. With a molecular weight over 80 MDa, each IFT train is a dynamic polymer of two large complexes (IFT-A and -B) and motor proteins, posing a formidable challenge to mechanistic understanding. Here, we reconstituted the complete human IFT-A complex and obtained its structure using cryo-EM. Combined with AlphaFold prediction and genome-editing studies, our results illuminate how IFT-A polymerizes; interacts with IFT-B; and uses an array of β-propeller and TPR domains to create “carriages” of the IFT train that engage TULP adaptor proteins. We show that IFT-A·TULP carriages are essential for cilia localization of diverse membrane proteins, as well as ICK – the key kinase regulating IFT train turnaround. These data establish a structural link between IFT-A’s distinct functions, provide a blueprint for the IFT-A train, and shed light on how IFT evolved from a proto-coatomer ancestor.

## INTRODUCTION

Intraflagellar transport (IFT) trains are multi-megadalton protein assemblies that traffic cargoes and signaling molecules into and out of cilia, the ubiquitous antenna-like organelles (Rosenbaum and Witman, 2002). Without IFT trains, cilia cannot assemble; sense or transduce signals; or generate the beating motions that power fluid flow and cell locomotion. Thus, IFT trains are essential for human life, and partial loss of function causes a range of human disorders (“ciliopathies”) associated with vision impairment, skeletal abnormalities, cystic kidneys, and infertility, among other conditions (Reiter and Leroux, 2017).

Consisting of ~1000 protein subunits, each IFT train is built around arrays of two large complexes (IFT-A and -B), of which IFT-A is the focus of this study (Figure 1A) (Cole et al., 1998; Jordan et al., 2018; Kozminski et al., 1993; Pigino et al., 2009; Piperno and Mead, 1997). Each train also incorporates two types of motor protein (kinesin-2 and dynein-2) and binds to cargoes, either directly or via associating factors (e.g. the BBSome and TULP family proteins) [reviewed in (Jordan and Pigino, 2021; Nachury, 2018; Webb et al., 2020)]. In cells, IFT trains and cargoes assemble at the base of the cilium; move through the ciliary diffusion barrier (the transition zone); then travel in the confined space between the cilium’s axoneme and membrane, driven by kinesin-2 (a process termed anterograde IFT) (van den Hoek et al., 2022; Kozminski et al., 1995; Wingfield et al., 2017). At the tip, IFT trains restructure, often releasing cargoes, and return to the cell body under the power of dynein-2 (retrograde IFT) (Lechtreck, 2015; Pazour et al., 1999; Porter et al., 1999; Signor et al., 1999; Stepanek and Pigino, 2016). The IFT train turnaround process is mysterious but is critically regulated by a tip-localized kinase named ICK/CILK1 in mammals (Broekhuis et al., 2014; Chaya et al., 2014; Nakamura et al., 2020; Oh et al., 2019; Paige Taylor et al., 2016). In the absence of high-resolution structural information, understanding of IFT train mechanism – and thus the molecular basis of ciliary trafficking – has been hampered.

**Figure 1.**
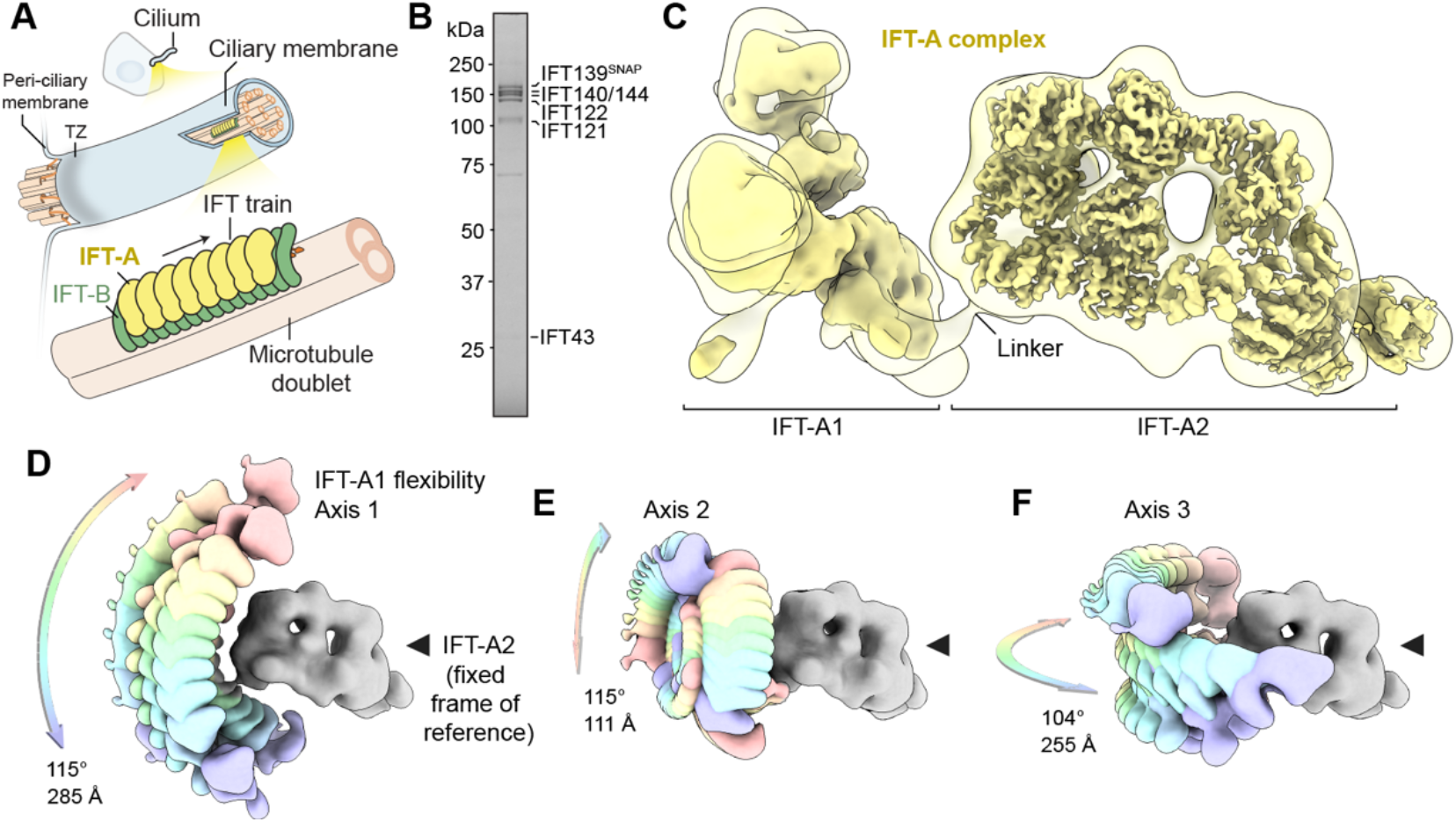
IFT-A Consists of Two Modules Connected by a Flexible Linker. (A) Schematic of an anterograde IFT train, consisting of polymeric IFT-A and IFT-B complexes, traveling along a microtubule doublet toward the tip of the cilium. Dynein-2 is omitted for clarity. TZ; transition zone. (B) SDS-PAGE of purified human IFT-A complex. See also Figure S1. (C) Cryo-EM structure of IFT-A complex, which consists of two modules (IFT-A1 and IFT-A2). Individual IFT-A1 and IFT-A2 maps are docked into a combined map (transparent envelope) from RELION multi-body analysis (Nakane et al., 2018) (see Methods). IFT-A2 map is locally sharpened using LocScale (Jakobi et al., 2017). IFT-A1 map is unsharpened. Flexible linker connecting the two modules indicated. See also Figure S1. (D – F) Different positions of IFT-A1 module (rainbow color scheme) with respect to IFT-A2 (gray; fixed frame of reference). Three principal components, representing rotations about three orthogonal axes, were determined using RELION multi-body analysis. The maximal rotation angle and displacement of the IFT-A2 module are indicated in each case.

Studies of similarly large and complex macromolecular machines, such as the nuclear pore complex, have been aided by the availability of structural models derived from docking atomic coordinates of subunits into more complete but lower resolution maps from cryoelectron tomography (cryo-ET) (Schwartz, 2022). Pioneering studies have produced cryo-ET maps of anterograde IFT trains *in situ* at 24–37 Å resolution (van den Hoek et al., 2022; Jordan et al., 2018); crystal structures of IFT-B sub-complexes (Bhogaraju et al., 2011, 2013; Taschner et al., 2014, 2016, 2018; Wachter et al., 2019); and the cryo-electron microscopy (cryo-EM) structure of the dynein-2 complex (Toropova et al., 2019). With the exception of dynein-2, however, the relatively small size and shape of solved IFT domains has made it impossible to dock them unambiguously into cryo-ET maps, and thus the location of individual proteins within the IFT train has been unknown.

IFT-A is a principal complex of the IFT train, first characterized in *C. reinhardtii* (Cole et al., 1998; Piperno and Mead, 1997). Each IFT-A complex has a mass of 768 kDa in humans and consists of six subunits. Biochemical studies have suggested that three of the IFT-A subunits (IFT144/IFT140/IFT122) form a “core”, whose disruption has a severe impact on cilia formation and trafficking (Behal et al., 2012; Hirano et al., 2017; Mukhopadhyay et al., 2010), whereas the others (IFT139/IFT121/IFT43) form a “non-core” sub-complex, which also serves important roles (reviewed in Taschner and Lorentzen, 2016). These labels may not reflect the physical organization of IFT-A, however, as no molecular resolution structures of the IFT-A complex nor any of its subunits have been reported. Strikingly, cryo-ET has revealed that the IFT-A complex lies directly underneath the ciliary membrane in anterograde IFT trains (Jordan et al., 2018), compatible with its role in the ciliary import of a variety of membrane proteins, including G-protein coupled receptors (GPCRs) (Fu et al., 2016; Hirano et al., 2017; Liem et al., 2012; Mukhopadhyay et al., 2010; Picariello et al., 2019). Moreover, sequence analysis has suggested that four of the IFT-A subunits (IFT144, IFT140, IFT122, and IFT121) are distantly related to coat proteins, in particular COPI/coatomer, which polymerizes on membranes (Avidor-Reiss et al., 2004; van Dam et al., 2013; Jékely and Arendt, 2006; Quidwai et al., 2021; Taschner et al., 2012). Conversely, foundational work and subsequent studies have shown that IFT-A mutants perturb retrograde IFT, leading to short cilia with accumulations of material at their tip (Iomini et al., 2001; Piperno et al., 1998). Thus, IFT-A appears to function in membrane protein transport into cilia as well as retrograde IFT, potentially as a dynein-2 activator (Pedersen et al., 2006; Williamson et al., 2012). How these different activities of IFT-A are coupled remains unclear.

Here, we reconstituted the human IFT-A complex and overcame its remarkable flexibility to obtain its cryo-EM structure up to 3.5 Å resolution. Our results reveal an unanticipated organization of subunits that could be docked unambiguously into previous cryo-ET maps, allowing us to build a pseudoatomic model of the IFT-A polymer and design structure-based mutants that we introduced into genome-edited mammalian cells. Our results illuminate how IFT-A polymerizes, interacts with IFT-B, and uses an array of β-propeller and TPR domains to create “carriages” of the IFT train that engage TULP adaptor proteins. We show that IFT-A·TULP carriages are not only required for cilia localization of diverse membrane proteins, but also ICK, the key kinase that regulates IFT train turnaround. These data establish a link between IFT-A’s distinct functions, provide a molecular picture of IFT-A in the train, and shed light on how IFT evolved from a proto-coatomer ancestor.

## RESULTS

### IFT-A Consists of Two Flexibly Connected Modules

To elucidate the structure and mechanism of human IFT-A, we affinity purified its components (IFT144·IFT140·IFT122; IFT121·IFT43; and IFT139), assembled them *in vitro*, and then isolated the complex using size-exclusion chromatography (Figure S1). This protocol yielded stoichiometric IFT-A (Figure 1B) that was amenable to analysis by cryo-EM (Figure 1C; Figure S1). Single-particle image processing revealed that IFT-A consists of two modules – which we term IFT-A1 and IFT-A2 – connected by a flexible linker (Figure 1C). IFT-A2 is a comparatively rigid structure, which we determined to an average resolution of 3.5 Å (Figure S1). IFT-A1 displays continuous flexibility, both internally and in its position relative to IFT-A2, and was resolved to 7–15 Å resolution (Figure S1). The IFT-A1 module flexes around three principal axes with respect to IFT-A2, moving by up to 115° and 285 Å about the flexible linker (Figure 1D–F; Movie S1). This striking flexibility may enable IFT-A to adopt the different curvatures observed for IFT trains queuing at the ciliary base and traveling along the ciliary shaft (van den Hoek et al., 2022; Jordan et al., 2018).

### Structure of the IFT-A2 Module

The well-resolved map of IFT-A2 enabled building of an atomic model (Figure 2; Table S1,S2; Methods). This revealed that IFT-A2 is composed of IFT139, IFT121, IFT43, and the N-terminal half of IFT122 (Figure 2A–C). Side-chain densities enabled unambiguous assignment of IFT122, IFT121, and IFT139 (Figure 2C,E) and a high-confidence AlphaFold (AF) Multimer prediction confirmed assignment of IFT43 (Figure S2B) (Evans et al., 2021). The composition of IFT-A2 is notable, as IFT-A2 can be viewed as the rigid core of IFT-A, yet previously IFT139, IFT121, and IFT43 have been termed non-core subunits. To avoid confusion, here we refer to IFT139, IFT121, and IFT43 as IFT-A2 subunits.

**Figure 2.**
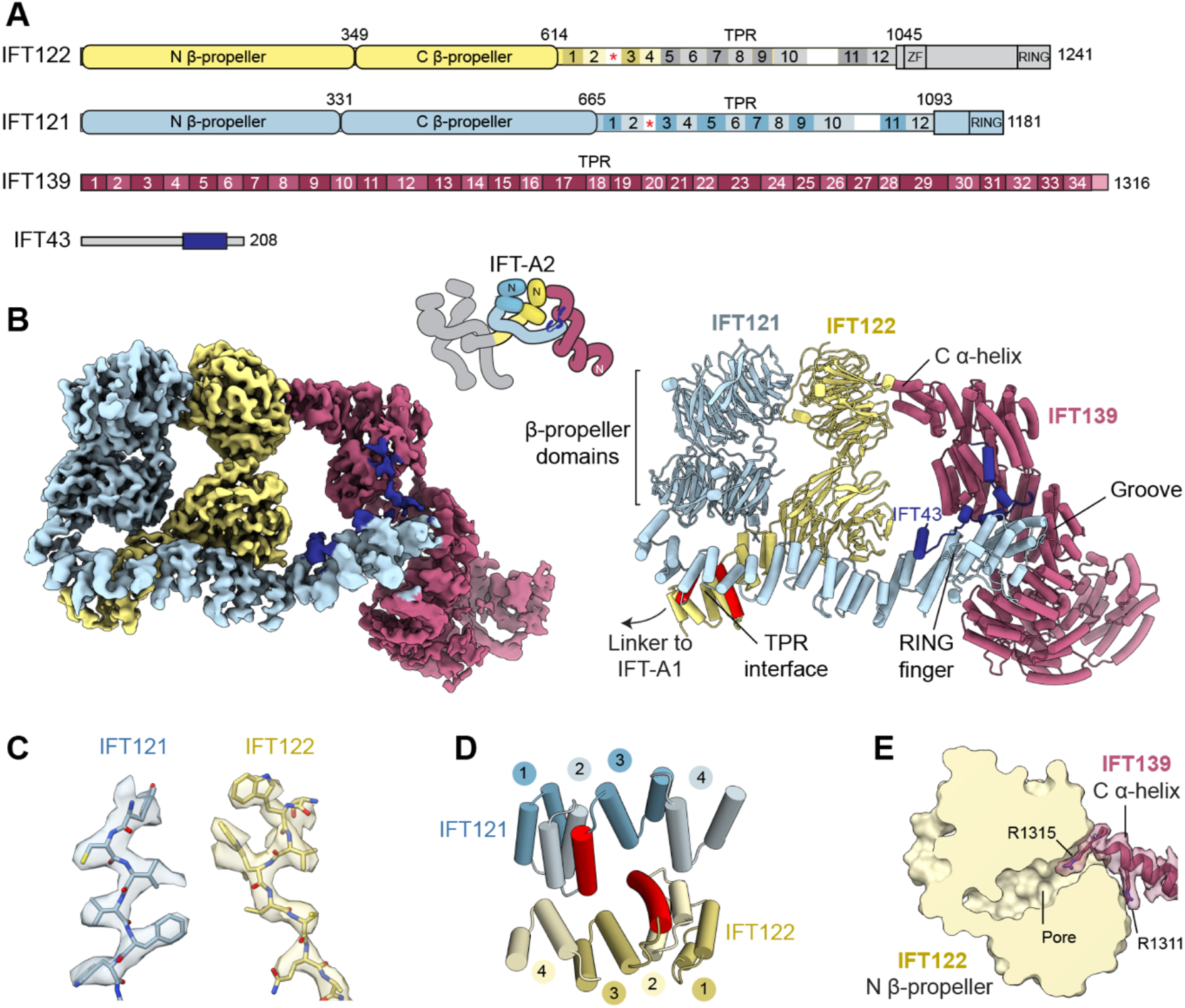
The IFT-A2 Module is an Interconnected Assembly of IFT122, IFT121, IFT139, and IFT43. (A) Domain organization of IFT-A2 module proteins. Red asterisk; α-helix insert. ZF; zinc finger domain. RING; RING finger domain. Numbering within domains – TPR repeats; outside – residues. IFT122 C-terminal region (gray) is in the IFT-A1 module. (B) Left, cryo-EM map of IFT-A2 locally sharpened using LocScale and colored by subunit as in (A). Inset, schematic cartoon, N-termini indicated. IFT-A1 in gray. Right, atomic model of IFT-A2 module. Key structural features are indicated. The distinctive single α-helix insert in IFT122 and IFT121 is colored red. (C) Example EM density and atomic coordinates for IFT121 and IFT122; side-chain resolution enables unambiguous protein assignment. (D) Enlarged view of the TPR interface between IFT121 and IFT122. The distinctive single α-helix insert in each subunit is colored red. TPR repeats are numbered. (E) Interface between IFT139 C α-helix and IFT122 β-propeller. EM density and atomic coordinates shown for IFT139. IFT122 is shown in surface representation. Cross-section view through β-propeller shown. Two well-resolved arginine residues (R1311 and R1315) involved in interaction are labeled.

IFT122 and IFT121 share a similar architecture (Figure 2A). Both consist of two β-propeller domains, followed by C-proximal tetratricopeptide repeats (TPRs). This domain layout is similar to the COPI subunits α and β’, confirming earlier predictions (Avidor-Reiss et al., 2004; van Dam et al., 2013; Jékely and Arendt, 2006; Quidwai et al., 2021; Taschner et al., 2012). Our structure shows that IFT121 also has a degenerate RING finger within its C-terminal domain (CTD) (Figure 2A,B), echoing COPI, HOPS, CORVET, and GATOR2 (Hunter et al., 2017; Kaur and Subramanian, 2015; Valenstein et al., 2022). In IFT121, the RING finger domain mediates interaction with IFT43 (Figure 2B) and mutations at this interface are associated with cranioectodermal dysplasia (see Figure S3 for mapping of these and other human disease-causing mutations in the complex, curated in Table S3).

Within the IFT-A2 module, IFT122 and IFT121 interact extensively through their β-propeller domains (Figure 2B) and via their TPR regions, which bind to each other with 2-fold pseudosymmetry (Figure 2D). The TPR interface involves a distinctive single α-helix inserted between the second and third TPR in each subunit (Figure 2D; red cylinders). We find that this α-helix insert is a signature motif of the IFT machinery, which recurs in IFT-A1, and also in IFT-B (see Discussion).

IFT122, specifically its TPR region, extends out of IFT-A2 and forms the flexible linker to the other module of IFT-A (IFT-A1) (Figure 1C). In contrast, the TPR region of IFT121 curves into IFT-A2, where it is anchored by interactions with IFT139 and IFT43 (Figure 2B).

IFT139 consists of a superhelix of 34 TPRs capped by a C-terminal α-helix, which inserts into the N β-propeller of IFT122 (Figure 2B,E). A groove in the IFT139 superhelix accommodates the C-terminal end of IFT121 (Figure 2B). The IFT-A2 module is completed by IFT43, whose conserved C-terminal region (Zhu et al., 2017) bridges IFT121 and IFT139 (Figure 2B). Together, this interconnected assembly gives IFT-A2 structural stability and suggests why the central IFT122 and IFT121 subunits are crucial for IFT-A integrity (Behal et al., 2012; Hirano et al., 2017; Mukhopadhyay et al., 2010).

### Structure of the IFT-A1 Module

Our cryo-EM map of the IFT-A1 module shows two pairs of β-propeller domains (Figure 3A,B), which are splayed apart. To establish which pair belongs to IFT144, we engineered an IFT-A complex lacking the IFT144 N β-propeller (IFT144ΔN) and imaged it by cryo-EM. With the structure oriented as in Figure 3C, the missing density assigns the β-propellers in the 12 o’clock position to IFT144. We then used AF Multimer to generate a pseudoatomic model of IFT144, IFT140, and the C-terminal region of IFT122 (Figure 3A,B). The ternary complex, which was predicted with high confidence (Figure S2), fits unambiguously into the IFT-A1 cryo-EM map and indicates that the C-terminal regions of IFT144 and IFT140 are flexible and unresolved (Figure 3B). The extensive interface between the IFT122 CTD and IFT144·IFT140 is in agreement with biochemical data identifying the IFT122 C-terminal region as essential for interaction with IFT144·IFT140 (Takahara et al., 2018). Strikingly, IFT144 and IFT140 interact via a TPR interface (Figure 3D) that is superimposable (RMSD 0.99 Å) with the one we found experimentally in IFT122 and IFT121 (including the distinctive α-helix insertion) (Figure 2D). These results indicate that IFT144·IFT140 and IFT122·IFT121 may have originated from a gene duplication event that was instrumental in the formation of the IFT-A complex.

**Figure 3.**
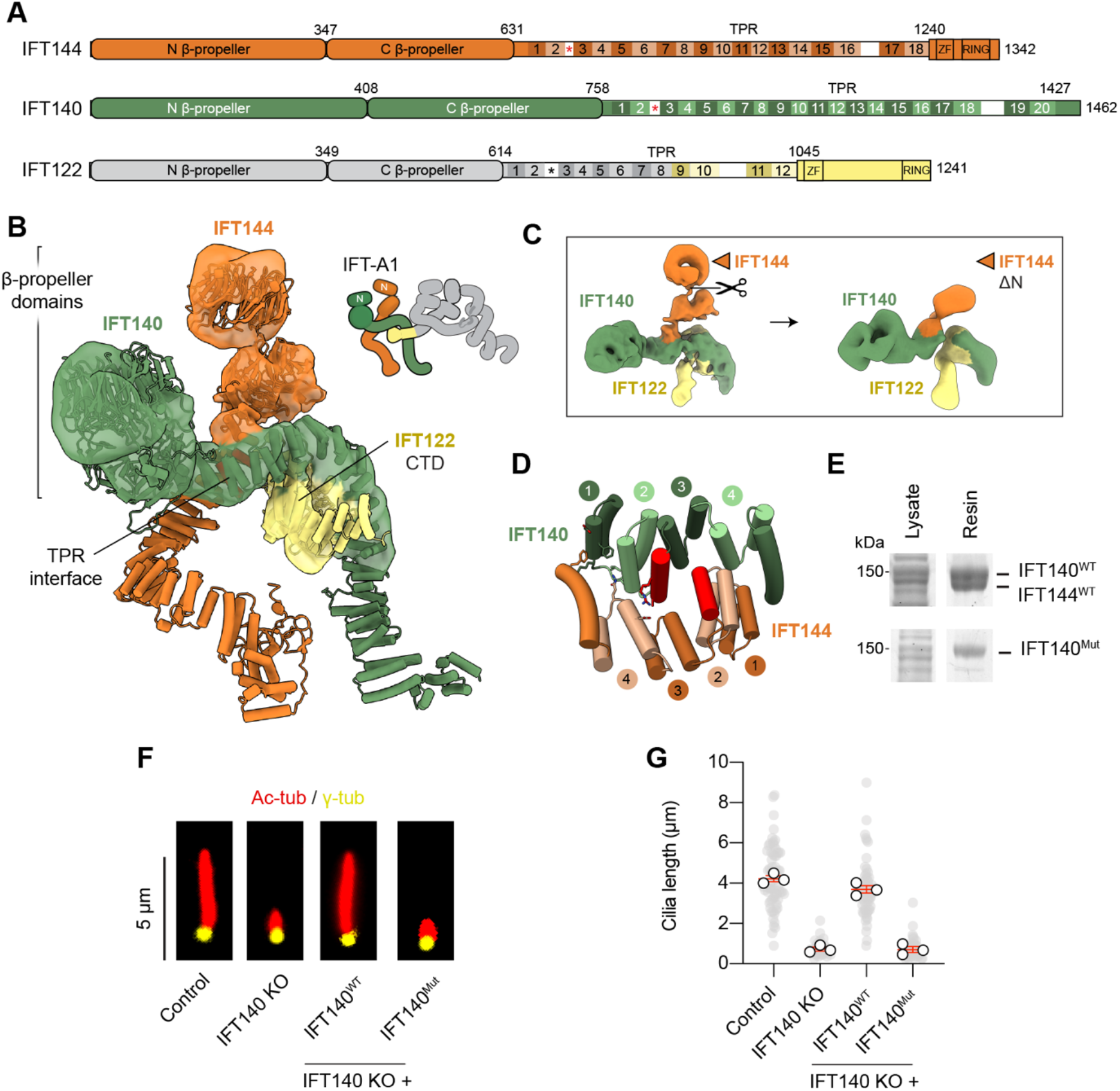
The IFT-A1 Module is Formed by IFT144, IFT140, and the C-terminal Domain of IFT122. (A) Domain organization of IFT-A1 module proteins. Red asterisk; α-helix insert. ZF; zinc finger domain. RING; RING finger domain. Numbering within domains – TPR repeats; outside – residues. IFT122 N-terminal region (gray) is in the IFT-A2 module. (B) Cryo-EM structure of IFT-A1 with AF Multimer model fitted, colored as in (A). Inset, schematic cartoon, N-termini indicated. See also Figure S2. (C) Comparison of IFT-A1 structures containing full-length IFT144 (left) and IFT144ΔN (right), in which the IFT144 N-terminal β-propeller was deleted. This missing density assigns the β-propellers in the 12 o’clock position to IFT144. (D) Close-up view of TPR interface between IFT140 and IFT144. Compare with Figure 2D. The distinctive single α-helix insert in each subunit is colored red. TPR repeats numbered. Residues in IFT140 chosen for mutagenesis are shown in stick representation. (E) SDS-PAGE of IFT144·IFT140 binding assay. Wild-type IFT140 co-purifies with IFT144 (upper panel), whereas IFT140^Mut^ with interface mutations does not (lower panel) (IFT140^Mut^: D789R/F792A/K796E/V822A/N826A/A830Y/A833Y/R837E). IFT140^Mut^ purification itself unaffected. (F) Representative images of IMCD-3 cilia fixed and immunofluorescently labeled with acetylated tubulin (red) and gamma tubulin (yellow) to identify cilia and basal bodies respectively. (G) Quantification of cilia length experiment shown in the indicated cells. Individual data points (gray circles), averages of separate experiments (white circles). Red lines; mean (± SEM). Control n = 70; IFT140 KO n = 33; IFT140 KO + IFT140^WT^ n = 65, + IFT140^Mut^ n = 44 cilia analyzed. One-way ANOVA followed by Kruskal-Wallis test values, control vs IFT140 KO *p* < 0.0001; control vs IFT140^WT^ *p* > 0.5; control vs IFT140^Mut^ *p* < 0.0001.

To test the predicted TPR interface between IFT144 and IFT140, we mutated conserved interface residues in IFT140 (Figure 3D). Whereas IFT144 co-purified robustly with wild-type IFT140 (Figure 3E; upper panel), this interaction was almost completely abolished by the IFT140 interface mutations (which had no effect on purification of IFT140 itself) (Figure 3E; lower panel). We next established the effect of the mutations on ciliogenesis in IMCD-3 cells; a widely used mouse kidney cell line that forms primary cilia upon serum starvation (cilia length of 4.2 ± 0.1 μm [mean ± SEM]) (Figure 3F,G). CRISPR knockout (KO) of IFT140 (Figure S4) reduced the cilia to short stumps (0.7 ± 0.1 μm) that barely extended beyond the basal body (Figure 3F,G). Stable expression of wild-type IFT140 restored the cilia length to 3.7 ± 0.2 μm; a value not significantly different to the control (*p* > 0.05). In contrast, the IFT140 interface mutant completely failed to rescue cilia length (0.7 ± 0.1 μm) (Figure 3F,G), despite being expressed at comparable levels to the wild-type construct (Figure S4F). These data establish that IFT144 and IFT140 interact via their TPR regions and that this interface is vital for proper cilia formation.

### Structure of the Complete IFT-A Complex

Combining our structures of IFT-A1 and -A2 reveals the complete architecture of the IFT-A complex (Figure 4A). This vividly shows the structural relationship between the two pairs of COPI-like subunits: IFT144·IFT140 in the IFT-A1 module and IFT122·IFT121 in the IFT-A2 module (Figure 4A). Their N β-propeller domains lie on the top face of the complex, with their TPR regions lying below. The TPR region of IFT122 forms the flexible linker between the two modules. Our cryo-EM structure is consistent with the reported biochemical interactions between subunits (Behal et al., 2012; Hirano et al., 2017; Ishida et al., 2021; Mukhopadhyay et al., 2010; Takahara et al., 2018). It is also compatible with our CRISPR knockouts in IMCD-3 cells (Figure S4), showing that IFT144, IFT140, IFT122, and IFT121 are each critical for the formation of normal length cilia, whereas knock-out of IFT139 – which is less integral in the assembly – has a milder but still significant effect (Figure 4B) [see also (Hirano et al., 2017; Scheidel and Blacque, 2018; Yi et al., 2017)]. In each case, stable expression of the missing IFT-A subunit restored cilia length to the level of the control, confirming specificity (Figure S4). These knockout cell lines provided a blank canvas in which structure-based mutants of the IFT-A subunits could be introduced to dissect their roles (described below).

**Figure 4.**
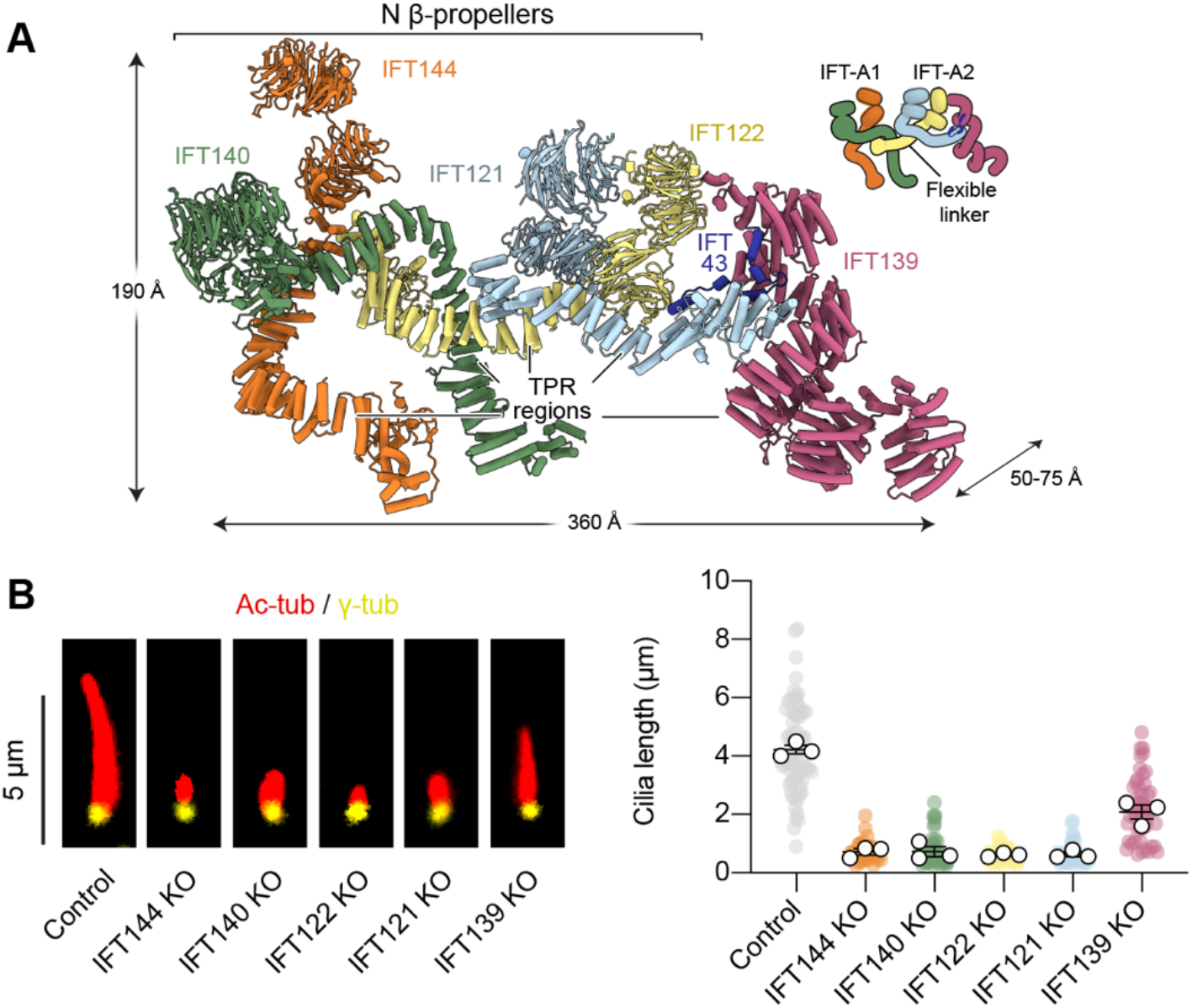
Architecture of the Complete IFT-A Complex and Importance of its Subunits in Cilia Formation. (A) Atomic model of complete IFT-A complex. Key structural features labeled and colored by subunit. Labeled arrows show the dimensions of the complex. Inset, cartoon schematic. The position of IFT-A1 and IFT-A2 represents the average conformer obtained from RELION multi-body analysis (Figure 1D–F). (B) Left, representative immunofluorescently labeled IMCD-3 cilia in control cells and IFT-A subunit knockout (KO) cells. Cells were fixed and immunofluorescently labeled with acetylated tubulin (red) and gamma tubulin (yellow) to identify cilia and basal bodies respectively. Right, quantification of cilia length from three separate experiments. Individual data points colored by subunit, control in gray. White circles; average from each separate experiment. Control cell data is repeated from Figure 1G for ease of comparison. Lines; mean (± SEM). Control n = 70; KOs; IFT144 n = 34, IFT140 n = 51, IFT122 n = 49, IFT121 n = 41, IFT139 n= 50 cilia analyzed. One-way ANOVA followed by Kruskal-Wallis test values, control vs IFT144 KO *p* < 0.0001; control vs IFT140 KO *p* < 0.0001; control vs IFT122 *p* < 0.0001; control vs IFT121 *p* < 0.0001; control vs IFT139 *p* < 0.01.

### IFT-A Polymerizes into “Carriages” with its β-propeller Domains Facing the Ciliary Membrane

Cryo-ET has revealed the morphology of the *C. reinhardtii* IFT-A polymer within anterograde IFT trains at 24–33 Å resolution *in situ* (van den Hoek et al., 2022; Jordan et al., 2018). To understand where individual proteins are located, we docked our atomic models of human IFT-A1 and -A2 into the cryo-ET map (EMDB-4304) (Jordan et al., 2018) (Figure 5A). A global fitting analysis showed that IFT-A2 module fits unambiguously into one side of the polymer (Figure 5B). IFT-A1 shows an unambiguous fit into the remaining density, following gentle bending motions within its flexible regions, which we modelled using molecular dynamics flexible fitting (Figure 5C; Movie S2). Our analysis assigns bulbous densities in the cryo-ET map to the β-propeller domains IFT144, IFT140, IFT122, and IFT121 (Figure 5D). The TPR regions of IFT122, IFT140, and IFT144 also display strong density, indicating that these flexible domains are held in place in the polymer (Figure 5D). To generate a model of the IFT-A train, we docked successive copies of the IFT-A complex (Figure 5E; Movie S2). For visualization purposes, 14 copies are displayed in Figure 5E, representing an IFT-A train of 11 MDa in mass and ~1,700 Å in length; about half the length of a typical anterograde train in *C. reinhardtii* (Jordan et al., 2018). Strikingly, each IFT-A complex is positioned so that the N β-propeller domains from IFT144, IFT140, IFT122 and IFT121 all lie next to the ciliary membrane (Figure 5E; circles). The IFT-A complexes in the polymer are notably interwoven. For example, the IFT140 β-propeller domains from one complex reach over and contact the IFT121 β-propellers from the adjacent complex (Figure 5F). Moreover, the TPR region of IFT140 from one complex wraps around the TPR region of IFT144 from the adjacent complex, forming an interlinked arrangement (Figure 5G). Together, the arrangement of β-propeller and TPR domains in the IFT-A polymer creates a series of large open compartments adjacent to the ciliary membrane (Figure 5E). We refer to these compartments as the “carriages” of the IFT train.

**Figure 5.**
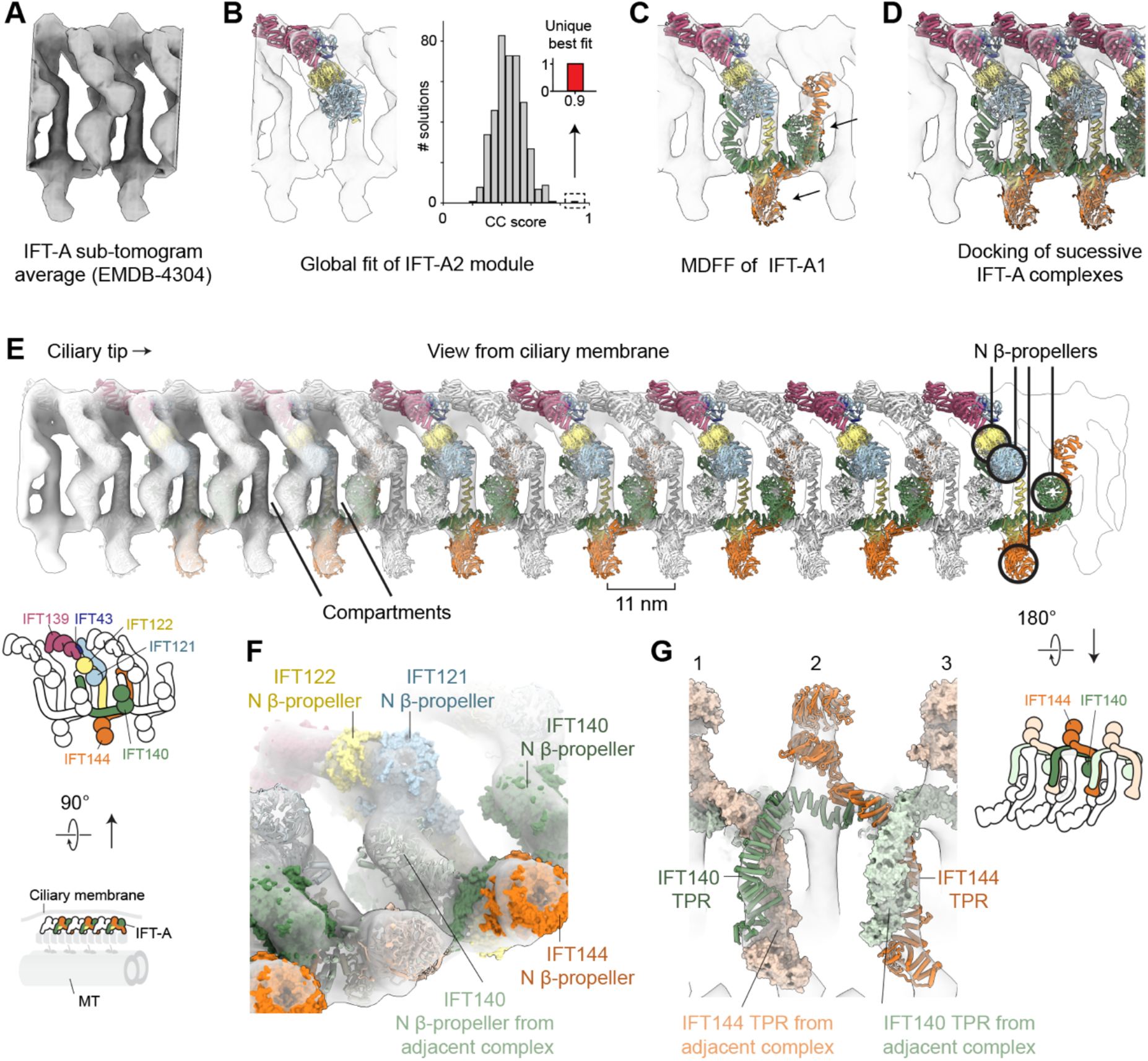
IFT-A Polymerizes into “Carriages” with its β-propeller Domains Facing the Ciliary Membrane. (A) Sub-tomogram average of IFT-A complex within anterograde IFT trains at 33 Å resolution (*C. reinhartdtii*, EMDB-4303) (Jordan et al., 2018). (B) Docking of the IFT-A2 module into the cryo-ET map using a global fitting procedure from 10,000 random starting positions. The unique top-scoring fit (depicted in the left-hand panel) has a high cross-correlation (CC) score (0.89; red bar, right-hand panel) that is clearly discriminated from the lower scoring false-positive distribution (gray bars). (C) IFT-A1 module fits with gentle bending of flexible regions (arrows) into EM density using molecular dynamics flexible fitting (MDFF). See also Movie S2. (D) Docking of successive IFT-A complexes to generate the IFT-A polymer. (E) Pseudo-atomic model of anterograde IFT-A polymer; 14 individual IFT-A complexes in alternating color/white representation for distinction. Cryo-ET map of IFT-A polymer (EMDB-4304) shown in gradient transparency, from left to right. View of train looking down from ciliary membrane. β-propellers of IFT144, IFT140, IFT122 and IFT121 surround large open compartments. N-terminal β-propellers (circled) are adjacent to the ciliary membrane. Inset, cartoon representations of IFT-A polymer showing subunit arrangement and orientation relative to microtubule (MT) doublet and membrane. (F) Oblique view of the IFT-A polymer, with IFT-A complexes shown alternately in surface and ribbon representation for distinction. The IFT140 N β-propeller domain from one complex (ribbon) reaches over to contact the IFT121 N β-propeller from the adjacent complex (surface). (G) Polymerization interface of IFT-A. View of train looking up from MT doublet (view rotation by 180° relative to panel E). IFT-A complexes shown alternately in saturated ribbon and pastel surface representation for distinction, with all subunits except IFT144 and IFT140 removed for clarity. The TPR region of IFT140 wraps round IFT144 TPR region of the adjacent complex, and vice versa. Cryo-ET map (EMDB-4304) shown as transparent surface.

### IFT88 C-terminal Loop Interacts with IFT144 Cleft for IFT-A Entry into Cilia

We next used our structure and functional assays to explore how IFT-A and IFT-B interact within the anterograde IFT train (Figure 6A). Docking our atomic model of IFT-A relative to a cryo-ET map of IFT-B and dynein-2 (Jordan et al., 2018) revealed that the C-terminal region of IFT144 extends down to reach IFT-B (Figure 6B). This is consistent with biochemical studies identifying the C-terminal region of IFT144 as critical for interaction with the IFT88 subunit of IFT-B (Ishida et al., 2021; Kobayashi et al., 2021). We generated a high-confidence AF model of the interface between IFT144 and IFT88 (Figure 6C). Negative controls replacing IFT144 with IFT140 or IFT122 confirmed the specificity of this prediction (Figure S2D). In this model, the C-terminal 14 residues of IFT88 bind as a loop in a cleft involving TPR17 of IFT144 (Figure 6C). Notably, the IFT144-binding motif of IFT88 lies at the end of an 82-residue disordered stretch that extends from the IFT88 core (Figure 6C). These results suggest that the C-terminal region of IFT88 forms a flexible tether between IFT-A and -B, explaining how it could accommodate the non-integral repeating distances of IFT-A and -B (11 and 6 nm) in the anterograde IFT train (Jordan et al., 2018).

**Figure 6.**
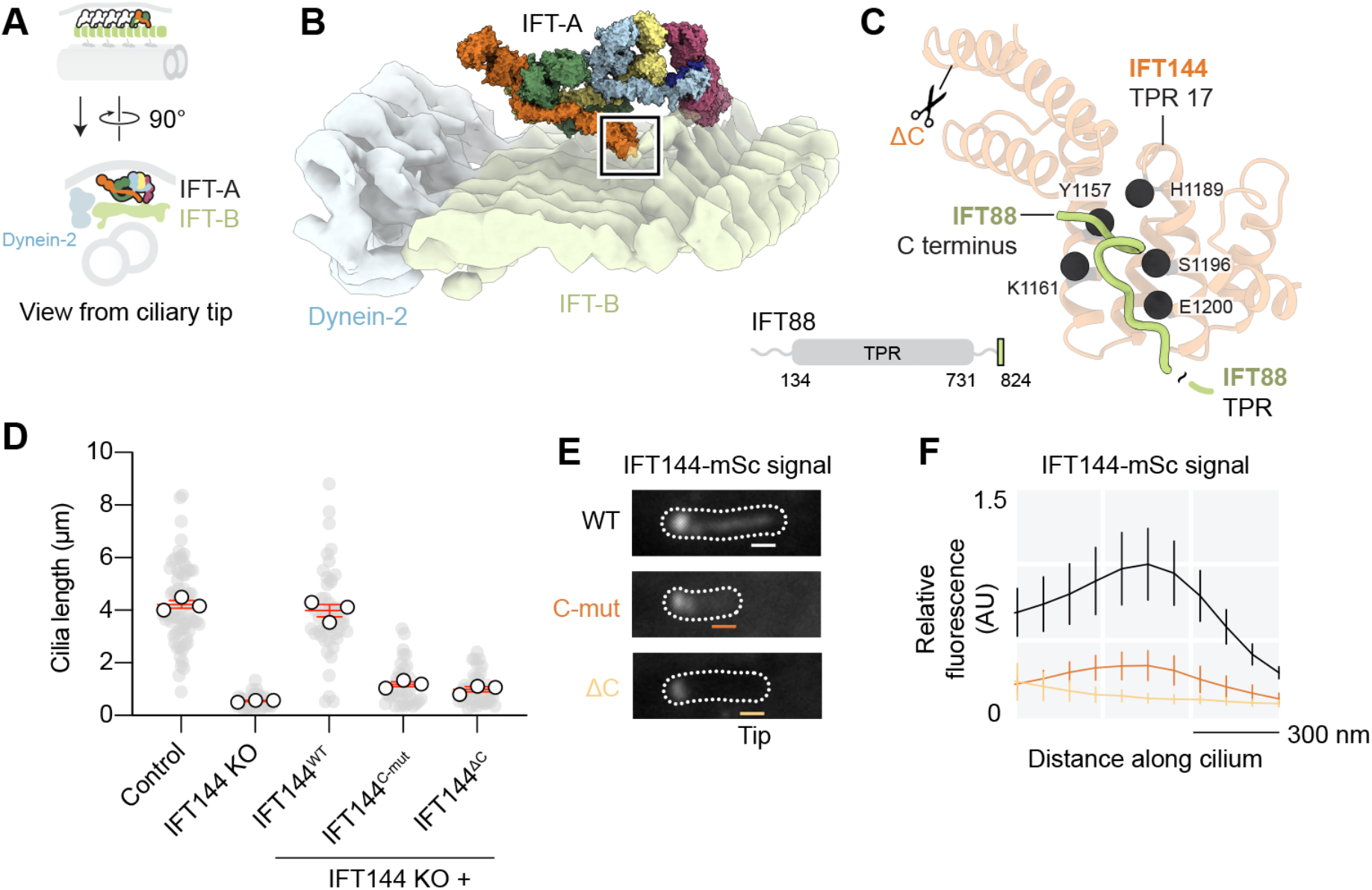
IFT88 C-terminal Loop Interacts with IFT144 Cleft for IFT-A Entry into Cilia. A) Top, schematic of anterograde IFT train with the IFT-A complex nearest the ciliary tip colored by subunit. Bottom, orthogonal view showing IFT-A relative to IFT-B, MT doublet, ciliary membrane and dynein-2. (B) Atomic model of IFT-A (surface representation, colored by subunit) shown relative to the sub-tomogram average of IFT-B and dynein-2 (EMDB-4303) (Jordan et al., 2018). The C-terminal region of IFT144 (boxed) reaches down to contact IFT-B. (C) Close-up of C-terminal IFT144 region (boxed in panel B), showing AF Multimer prediction of the interface between the IFT144 cleft and the C-terminal loop IFT88. IFT144 residues at the interface selected for mutagenesis in IFT144^C-mut^ (Y1157A/K1161E/H1189A/S1196A/E1200K) are depicted with spheres. Scissors represent site of IFT144 C-terminal truncation (residue 1103) in IFT144^ΔC^. Inset, IFT88 domain map, showing the C-terminal loop (green) at the end of an unstructured linker. (D) Quantification of cilia length in the indicated cells. Short-cilia phenotype in IFT144 KO cells cannot be rescued by stable expression of either IFT144^C-mut^ or IFT144^ΔC^ but is restored by expression of wild-type IFT144. Individual data points (gray circles) and averages of separate experiments (white circles) are shown. Red lines; mean (± SEM). Control cell data from Figure 3G displayed for comparison. Control n = 70; IFT144 KO n = 31; IFT144 KO + IFT144^WT^ n = 46, + IFT144^C-mut^ n =33, + IFT144^ΔC^ n = 41 cilia from three separate experiments. One-way ANOVA followed by Kruskal-Wallis test values, control vs IFT144 KO *p* < 0.0001; control vs IFT140^WT^ *p* > 0.5; control vs IFT144^C-mut^ *p* < 0.0001; control vs IFT144^ΔC^ *p* < 0.0001. (E) Representative images of mScarlet (mSc) tagged IFT144 constructs expressed in IFT144 KO cells. Cells also stably express mNG-DYNC2H1, which was used to mark cilia. Both IFT144^C-mut^ or IFT144^ΔC^ can be recruited to the ciliary base but are severely reduced within the cilium. For quantification in panel F, line-scans of IFT144-mSc fluorescence were generated near the ciliary tip to prevent overlapping signal from the base; bars are 900 nm representing the distance analyzed. (F) Plot of average IFT144-mSc fluorescent signal from line scans along the cilium length, aligned at the ciliary tip (peak in the IFT144^WT^ black trace). Values are normalized relative to IFT144^WT^ peak value. Traces show mean intensity ± SEM; n = 18 (WT), 8 (IFT144^C-mut^), 7 (IFT144^ΔC^) cilia measured from three separate experiments.

To test the importance of the predicted interface between IFT144 and IFT88, we mutated conserved interface residues in IFT144 (Figure 6C) and expressed the construct in IFT144 KO cells (Figure 6D–F). We compared this mutant (IFT144^C-mut^) to wild-type IFT144 and a characterized IFT144 C-terminal truncation (IFT144^ΔC^), which is associated with cranioectodermal dysplasia (Bredrup et al., 2011) and disrupts interaction between IFT-A and -B (Ishida et al., 2021). Stable expression of IFT144^C-mut^ or IFT144^ΔC^ failed to rescue proper cilia formation (Figure 6D). Moreover, while IFT144^C-mut^ could be recruited to the cilia base, its levels within cilia were drastically reduced compared to wild-type IFT144 (Figure 6E,F). This behavior mirrored that of the IFT144^ΔC^ construct (Figure 6E,F), which cannot effectively bind IFT-B (Ishida et al., 2021), and the impact of an IFT88 mutant lacking the C-terminal 28 residues (Ishida et al., 2021; Kobayashi et al., 2021). These data do not rule out additional contacts between IFT-A and -B involving globular domains, which are evident in the cryo-ET map (Jordan et al., 2018) (Figure 6B). However, they demonstrate that the interaction between the C-terminal loop of IFT88 and cleft of IFT144 is functionally crucial, as its disruption leaves IFT-A stranded at the base of the cilium.

### IFT-A·TULP Carriages Localize Diverse Ciliary Membrane Proteins and the Key Kinase for Train Turnaround

Finally, we used our structure of IFT-A to dissect its function in membrane protein import into cilia and retrograde transport to the cell body. TULP adaptor proteins (TUBBY, TULP2, and the ubiquitously expressed TULP3) are key players in IFT-A-mediated membrane protein transport (Mukhopadhyay et al., 2010). We used AF to generate a high-confidence prediction of the interface between TULP3 and IFT-A and mapped it onto our IFT-A structure (Figure 7A,B). TULP3 residues 10–44 form an α-helix (TULP3 N helix) that binds in a channel in the CTD of IFT122, while also making a minor contact with IFT140 (Figure 7A). The interface involves an acidic patch on the IFT122 RING finger interacting with basic residues in the TULP3 N helix. The AF model is in excellent agreement with the TULP3 residues required for IFT-A binding (Figure 7C) (Mukhopadhyay et al., 2010) and IFT-A·TULP3 binding involving IFT122, IFT140, and IFT144 (while indicating that IFT144 has a mainly allosteric role) (Hirano et al., 2017; Mukhopadhyay et al., 2010).

**Figure 7.**
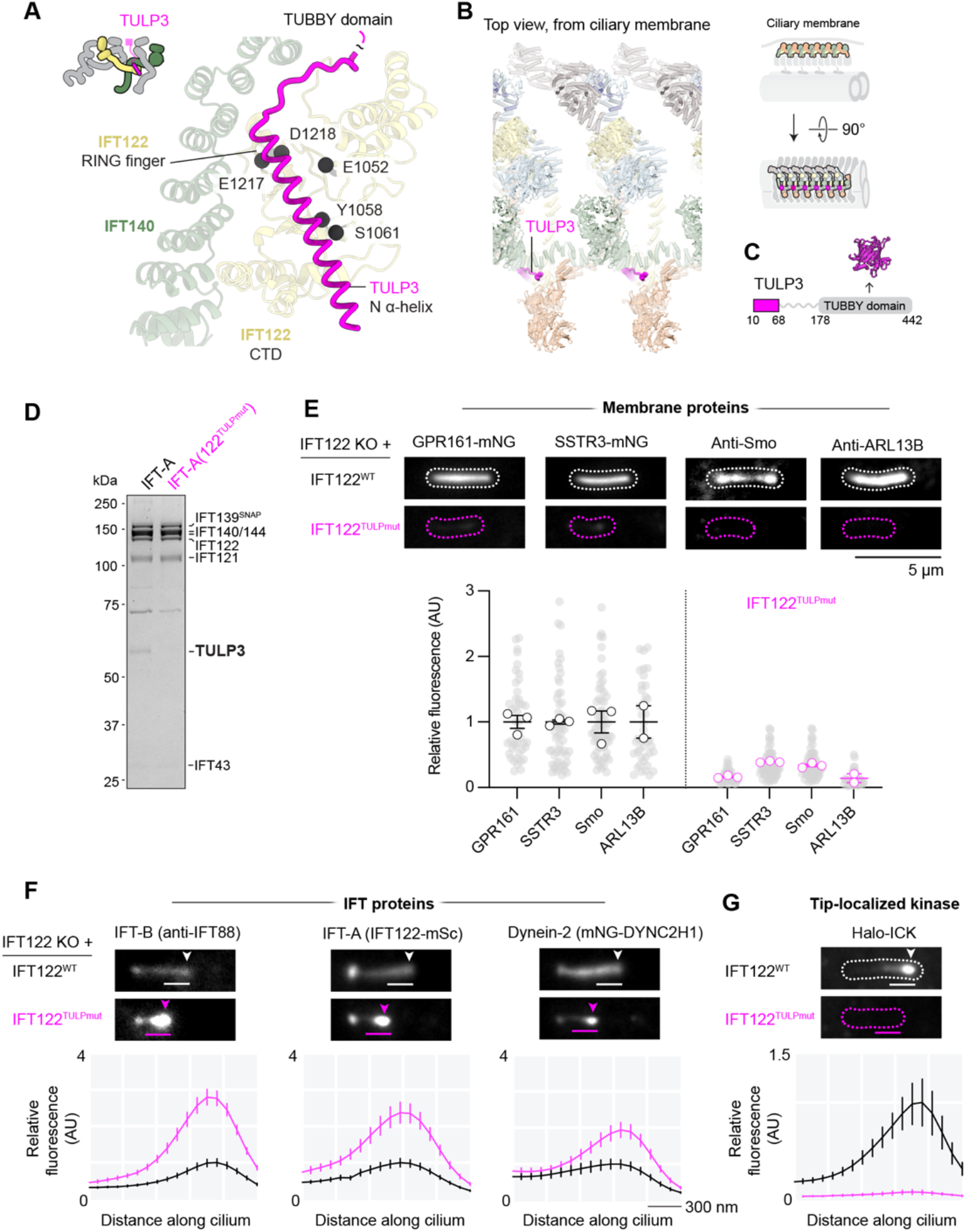
IFT-A·TULP Carriages Localize Diverse Ciliary Membrane Proteins and the Key Kinase for Train Turnaround. (A) AF model of the interface between TULP3 N-helix, IFT122 C-terminal domain (CTD) and IFT140. IFT122 residues at TULP interface selected for mutagenesis in IFT122^TULPmut^ (E1052R/Y1058A/S1062A/E1217R/D1218R) depicted with spheres. (B) Top view looking down from ciliary membrane, in the context of the anterograde IFT train (see schematic to the right). The TULP3 binding site gives the TUBBY domain direct access to the ciliary membrane. (C) Domain organization of TULP3, showing the TUBBY domain connected to the IFT-A binding N-helix by an unstructured linker. Crystal structure of TUBBY domain (PDB 1I7E; (Santagata et al., 2001)) shown at same scale as IFT-A polymer in (B). (D) Size-exclusion chromatography followed by SDS-PAGE of IFT-A and TULP3 (left) and IFT-A(122^TULPmut^) and TULP3 (right). TULP3 co-elutes with IFT-A but not IFT-A(122^TULPmut^). (E) Top, live-cell (GPR161-mNG and SSTR3-mNG) and immunofluorescence (Smo and ARL13B) images of membrane proteins in IFT122 KO cells stably expressing either wild-type IFT122 or IFT122^TULPmut^. From Smo experiments, cells were treated with 400 nM Smo agonist (SAG) to induce Smo ciliary localization. Bottom, quantification of average ciliary fluorescence, with IFT122^TULPmut^ values normalized relative to IFT122^WT^. Individual data points (gray circles), averages of separate experiments (white circles). Lines; mean (± SEM). IFT122^WT^ n = 44, 55, 64, 43, IFT122^TULPmut^ n = 46, 60, 43, 32 cilia analyzed for GPR161, SSTR3, Smo and ARL13B respectively, from 2–3 separate experiments. IFT122^WT^ vs IFT122^TULPmut^ Mann-Whitney test, *p* < 0.0001 for GPR161, SSTR3, Smo and ARL13B. (F) Top, immunofluorescence (IFT88) and live-cell (IFT122-mSc and mNG-DYNC2H1) images in IFT122 KO cells stably expressing either IFT122^WT^ or IFT122^TULPmut^. Bars are 1500 nm representing distance analyzed in lines scans below. Bottom panels, plots of average fluorescence intensity from line scans along the cilium length, aligned at the ciliary tip (peaks). Values are normalized relative to IFT122^WT^ peak values. Traces show mean intensity ± SEM; IFT122^WT^ n = 91, 59, 56, IFT122^TULPmut^ n = 70, 40, 31 cilia analyzed for IFT88, IFT122 and DYNC2H1 respectively, from three separate experiments. (G) Equivalent analysis for tip-localized kinase, ICK. Cilia (dashed outlines) marked by IFT122-mSc signal. Scale of x-axis same as panel F. IFT122^WT^ n = 41, IFT122^TULPmut^ n = 40 cilia analyzed, from three separate experiments.

To test if the IFT122 CTD is the crucial site for IFT-A·TULP3 association, we purified fulllength human TULP3 (Figure S6) and performed a size-exclusion-chromatography binding assay. Whereas TULP3 co-eluted with the wild-type IFT-A complex (Figure 7D, left-hand lane), binding was abolished by mutation of conserved interface residues in IFT122 CTD (IFT122^TULPmut^; which had no impact on IFT122 incorporation into the IFT-A complex; Figure 7D, right-hand lane). TULP2 and TUBBY were predicted to bind via the same interface, highlighting the IFT122 CTD as the conserved interaction site for TULP proteins (Figure S6). Downstream of the N helix, TULP proteins have an unstructured linker of ~100 amino acids, followed by a C-terminal TUBBY domain that is involved in phosphoinositide and membrane-protein binding (Figure 7C) (Badgandi et al., 2017; Mukhopadhyay et al., 2010; Santagata et al., 2001). Strikingly, in the IFT-A polymer, the TULP N helix binds in the large, open compartment that faces the ciliary membrane (Figure 7B), giving the TUBBY domain unfettered access to the lipid bilayer and membrane proteins.

Our structure-based IFT122 mutant (IFT122^TULPmut^) provides a unique tool to dissect IFTA mechanism, as it disrupts the TULP interface while leaving the other elements of IFT-A intact. Expression of IFT122^TULPmut^ in IFT122 KO IMCD-3 cells supported formation of cilia half the typical length (2.1 ± 0.03 μm) that were remarkably devoid of multiple classes of membrane protein (Figure 7E). Cilia localization of the GPCRs GPR161 and SSTR3 was virtually abolished, in line with their dependency on TULP3 for targeting (Mukhopadhyay et al., 2010, 2013). Agonist-induced cilia localization of Smoothened (Smo) was also virtually abolished (Figure 7E); a notable result given that Smo targeting has been found to be independent of TULP3 (Palicharla et al., 2021). This may reflect IFT122^TULPmut^ blocking binding to all TULP proteins rather than just TULP3. The small GTPase ARL13B – which has been placed at the top of the ciliary membrane protein hierarchy (Nachury and Mick, 2019) – also strictly depended on the IFT-A·TULP interface for cilia localization (Figure 7E) (Han et al., 2019; Palicharla et al., 2021). These data indicate that IFT-A·TULP carriages are master regulators of the ciliary membrane proteome.

Foundational studies implicated IFT-A in retrograde transport to the cell body rather than trafficking into cilia (Iomini et al., 2001; Piperno et al., 1998). To investigate if there is a connection between these activities, we quantified the localization of the IFT machinery in IFT122^TULPmut^ cells (Figure 7F). This showed that markers of IFT-B (IFT88), IFT-A (IFT122^TULPmut^), and dynein-2 (the motor subunit DYNC2H1) were each accumulated several-fold at the ciliary tip compared to control cells expressing wild-type IFT122 (Figure 7F), suggesting that the IFT-A·TULP interface is critical for proper retrograde IFT. The accumulation of both IFT-A and dynein-2 components at the ciliary tip is at odds with a simple model in which the presence of IFT-A is sufficient to activate dynein-2. This led us to hypothesize that another factor required for train turnaround could be missing. We therefore examined the localization of the kinase ICK; a central regulator of the transition between anterograde and retrograde IFT (Broekhuis et al., 2014; Chaya et al., 2014; Nakamura et al., 2020; Oh et al., 2019; Paige Taylor et al., 2016). Whereas ICK showed canonical tip-localization in control cells, its cilia localization was abolished in IFT122^TULPmut^ cells (Figure 7G). Thus, the IFT-A·TULP interface is essential for cilia localization of the key kinase regulating IFT train turnaround. These data establish a structural link between IFT-A’s roles in retrograde IFT and trafficking into cilia.

## DISCUSSION

IFT trains are molecular machines that regulate the proteome of cilia and flagella, underpinning the many sensory and propulsive functions of these organelles. Here we have combined cryo-EM, AF, structure-guided mutagenesis, biochemistry and cell biology to determine the structure of IFT-A and dissect its function. These data allow us to discuss fundamental questions concerning the mechanism of ciliary trafficking from a new structural perspective.

### Structural Organization of the IFT-A Complex

Our cryo-EM maps show that IFT-A is built around a dimer-of-dimers. The dimeric units (IFT144·IFT140 and IFT122·IFT121) form the basis for the two modules of IFT-A, which we call IFT-A1 and IFT-A2 [following the naming convention for the IFT-B subcomplexes, IFT-B1 and -B2 (Taschner et al., 2016)]. The architecture differs from two contemporaneous structural models of IFT-A based on fitting AF models into 18 Å and 23 Å resolution cryo-ET maps of the IFT-A polymer, recently deposited to bioRxiv (Lacey et al., 2022; McCafferty et al., 2022, respectively). Our high-resolution structure is in closer agreement with the preprint of Lacey et al., especially in the IFT-A2 module. The main difference is that, in the study of Lacey et al., IFT122-TPR is modelled as an isolated projection from IFT-A2, rather than a linker to IFT-A1 within the same complex. This may be due to ambiguity in how the modules are connected at 18 Å resolution (Lacey et al., 2022). Our structure of IFT-A is consistent with published interaction data (Hirano et al., 2017; Takahara et al., 2018) showing that the IFT122 C-terminal region forms the linker between IFT-A1 and IFT-A2.

### Evolution of IFT-A and IFT-B

The IFT144·IFT140 and IFT122·IFT121 subunits of IFT-A are related to the COPI subunits α and β’ (Quidwai et al., 2021), which also form a heterodimer (Lee and Goldberg, 2010). Our data reveal that the mode of dimerization in the IFT-A proteins is unique, involving a distinctive single α-helix inserted between the second and third TPR of each subunit. Interestingly, the other major complex of the IFT train – IFT-B – also contains a pair of β-propeller/TPR subunits (IFT172 and IFT80), which interact (Katoh et al., 2016). We used AF to generate a high-confidence prediction of the IFT172·IFT80 interface, which revealed the same signature α-helix inserted between the second and third TPR of each subunit (Figure S5). Together, these data suggest a model in which an ancestral COPI-like subunit evolved a dimerization interface by the acquisition of an extra α-helix between its second and third TPR, and this subunit was successively duplicated to form heterodimeric units of the IFT machinery: IFT172·IFT80, IFT144·IFT140, and IFT122·IFT121. In IFT-A, our data suggest that the addition of IFT139 and IFT43 to the complex had a critical role in rigidifying the IFT-A2 module, while leaving IFT-A1 flexible; a property that may help the IFT-A1 domains wrap around each other during IFT-A polymerization.

### Relationship Between IFT trains, Membrane Coat Proteins, and the Nuclear Pore Complex

At first sight, the organization of the IFT-A polymer appears remarkably different to its distant relatives COPI, COPII, clathrin, and the nuclear pore complex (NPC) (Beck et al., 2018). Whereas IFT-A forms an interwoven linear polymer in the anterograde IFT train, its closest relative, COPI, forms relatively discrete assembly units that coat vesicles (Dodonova et al., 2015, 2017). However, upon close inspection there are parallels. Our data show that the N β-propeller domains of IFT-A are all positioned adjacent the ciliary membrane. This mirrors the situation in COPI and the NPC, in which β-propeller domains are key membrane binding elements (von Appen et al., 2015; Dodonova et al., 2015), providing strong support for the proto-coatomer hypothesis (Devos et al., 2004). Moreover, while IFT-A polymerizes in a different fashion to COPI, the wrapping of IFT144·IFT140 TPR regions around each other in the IFT train is reminiscent of the way clathrin triskelia interdigitate to form cages (Fotin et al., 2004). These insights demonstrate the versatility of β-propeller/TPR subunits to form distinct polymeric architectures, while highlighting a common functional thread of selectively incorporating cargo into organelles.

### IFT-A β-propellers and TULP proteins in Membrane and Cargo Binding

Our structure shows how IFT-A uses its β-propeller and TPR domains to create a series of compartments that face the ciliary membrane in the anterograde IFT train. We show that TULP adaptor proteins bind in these carriages via the CTD of IFT122 and that this interface is essential for localization of several classes of membrane protein to cilia (GPR161 and SSTR3, rhodopsin-like GPCRs; Smo, a class-F GPCR; and ARL13B, a palmitoylated ARF-like GTPase). Our work targeting the TULP-binding interface on IFT-A is complementary to previous studies targeting TULP3 (Badgandi et al., 2017; Han et al., 2019; Mukhopadhyay et al., 2010, 2013; Palicharla et al., 2021). Structurally, the IFT-A carriages provide TULP proteins with direct access to the lipid bilayer, where the TUBBY domain is likely to interact with phosphatidylinositol 4,5-bisphosphate (PI(4,5)P_2_) and the ciliary targeting sequences on cargoes (Badgandi et al., 2017; Mukhopadhyay et al., 2010; Santagata et al., 2001), while being surrounded by IFT-A’s N β-propellers.

As the IFT-A N β-propellers and C-terminal region of IFT139 lie adjacent to the ciliary membrane, they may also contact the lipid bilayer directly. Indeed, an IFT-A sub-complex (IFT139·IFT121·IFT43) has been shown to associate with lipids and vesicles *in vitro*, with specificity for phosphatidic acid (Quidwai et al., 2021). In COPI, membrane-associated β-propeller domains interact with the dilysine motifs on cargoes destined for transport from the Golgi to the ER (Jackson et al., 2012; Ma and Goldberg, 2013). The sequence determinants for ciliary membrane protein targeting are less well defined, but can involve distributed, partially redundant motifs [for example, in the third intracellular loop and C-terminal region of ciliary GPCRs; (Barbeito and Garcia-Gonzalo, 2021)]. It is possible that the β-propellers of IFT-A and TUBBY domain of TULP work in conjunction to recognize different targeting elements on ciliary membrane proteins. Division of labor between IFT-A and TULP proteins may help explain why some organisms appear to lack IFT-A-binding TULP homologs yet are still able to concentrate a rich assortment of signaling receptors in their cilia (Sigg et al., 2017). Consistent with the idea of collaboration between IFT-A β-propellers and TUBBY domains, TULP4 – a divergent TULP family member – combines a TUBBY domain and a IFT121-like β-propeller in the same polypeptide (Mukhopadhyay and Jackson, 2011), which AF predicts to interact directly (Figure S6F).

### IFT-A Polymerization and Entry into Cilia

Where in the cell does IFT-A first polymerize and associate with cargoes? We did not observe spontaneous polymerization of IFT-A in solution, suggesting that IFT-A polymers may be stabilized by binding to IFT-B or the membrane. Striking views of assembling IFT trains show strings of IFT-B polymers extending from the cilia base into the cytoplasm, which could form a scaffold for IFT-A self-assembly (van den Hoek et al., 2022). In these flexible assembling trains, IFT-A carriages would be exposed on the surface, where they could latch onto cargoes in the nearby peri-ciliary membrane, which is a hotspot for vesicular fusion (Nachury et al., 2010). There is also evidence that IFT-A can associate with vesicles prior to their fusion with the peri-ciliary membrane (Quidwai et al., 2021). Here, the remarkable flexibility of IFT-A we describe may enable IFT-A to adapt to a spherical membrane. In either scenario, recruitment of IFT-A to membranes is likely to be regulated by exogenous factors, such as TULPs (Badgandi et al., 2017; Mukhopadhyay et al., 2010), CPLANE (Langousis et al., 2022; Toriyama et al., 2016), and small GTPases (Dewees et al., 2022; Langousis et al., 2022).

Once IFT-A is coupled to IFT-B and train assembly is complete, kinesin-2 would move the IFT train through the ciliary diffusion barrier (the transition zone) and into the cilium (van den Hoek et al., 2022; Lechtreck, 2015). Membranous cargoes would be dragged laterally from the peri-ciliary membrane through the transition zone (Figure 1A), which has both transmembrane and soluble components (Garcia-Gonzalo and Reiter, 2017). Here, the IFT-A·TULP carriages may help to displace transition zone proteins or ordered lipids that prevent free diffusion of membrane proteins into cilia. A deeper understanding of the physical nature of the transition zone will help to explore these possibilities.

### Role of IFT-A in Retrograde IFT

Our data shine new light on the IFT turnaround process at the ciliary tip, in which anterograde IFT trains convert into retrograde trains that travel back to the cell body under the power of dynein-2. Since the first characterized IFT-A mutants, it has been evident that partial loss of IFT-A function results in short cilia with bulbous accumulations at their tips, suggestive of defective retrograde transport (Iomini et al., 2001; Piperno et al., 1998). Mutations in dynein-2 cause similar accumulations (Pazour et al., 1999; Porter et al., 1999; Signor et al., 1999). More recently, it has been demonstrated that loss of a kinase, ICK, also results in aberrant tip accumulations, implicating ICK and its orthologs as a critical regulator of train turnaround (Broekhuis et al., 2014; Chaya et al., 2014; Nakamura et al., 2020; Oh et al., 2019; Paige Taylor et al., 2016). Clinically, mutations in IFT-A, dynein-2, and ICK are associated with a shared human disorder, short rib-polydactyly (Mitchison and Valente, 2017).

Here we show that when the IFT-A·TULP interface is disabled, targeting of ICK to the ciliary tip is abolished. Concurrently, IFT-A, IFT-B, and dynein-2 accumulate several-fold at the tip. This accumulation of IFT material closely resembles the impact of knocking out ICK (Nakamura et al., 2020). A notable difference is that ICK KO cells exhibit elongated cilia (Nakamura et al., 2020), likely because they over accumulate membrane proteins [a known driver of cilia extension (Ye et al., 2018)], whereas IFT122^TULPmut^ cells do not. Together, these data establish a link between the role of IFT-A in trafficking proteins into cilia and retrograde IFT.

Several new questions arise from this connection. First, is ICK, a kinase, transported by IFT-A directly? This would be surprising, given IFT-A’s specialization for membrane protein trafficking. Indeed, evidence suggests that ICK undergoes anterograde IFT by interacting directly with IFT-B (Nakamura et al., 2020). Thus, the essential role of IFT-A may be in trafficking a regulatory factor that tethers or releases ICK at the tip. Second, is the role of IFT-A in controlling kinase tip-localization conserved? Notably, global analysis of a destabilizing IFT140 mutant in *C. reinhardtii* showed that the abundance of several kinases was reduced (Picariello et al., 2019), including CDPK1 and mitogen-activated kinases related to ICK. These data point toward a conserved mechanism. In *C. reinhardtii*, IFT train turnaround can be evoked by applying pressure midway along the length of cilia (Nievergelt et al., 2022). As ICK undergoes anterograde IFT (Broekhuis et al., 2014; Nakamura et al., 2020), it will be interesting to examine if it accumulates at these pressure points. Finally, what are the targets of ICK that drive the turnaround process? To date, the major reported target of ICK and CDPK1 is kinesin-2 (Chaya et al., 2014; Liang et al., 2014), which is proposed to dissociate from the IFT train upon phosphorylation (Liang et al., 2014). Kinesin-2 dissociation may initiate the major conformational change that IFT trains undergo as they convert to retrograde trains (Chien et al., 2017; Jordan et al., 2018; Stepanek and Pigino, 2016; Wingfield et al., 2021; Zhang et al., 2021).

## Conclusion

Our structural and functional studies of the human IFT-A complex and pseudo-atomic model of the IFT-A train provide a foundation for understanding the mechanism of cilia trafficking and analysing patient mutations. From a technical standpoint, our work represents an integration of cryo-EM with AF predictions, which we validated using biochemistry and cell biology. The molecular picture of IFT-A that emerges from these studies should serve as a valuable tool to dissect how IFT-A re-organizes in the retrograde IFT train and how IFT-A·TULP carriages recognize targeting sequences on diverse transmembrane receptors, enabling cilia to function as cellular antennae that sense varied extracellular signals.

## ACKNOWLEDGEMENTS

We thank Peter Jackson (Stanford University) for IMCD-3 FlpIn cells; David J. Stephens (University of Bristol) for anti-IFT88; Luke Lavis (Janelia Research Campus) for JF646-HaloTag ligand; Andrew P. Carter and Chris Van Horn (MRC-LMB) for valuable discussions; Natasha Lukoyanova and Shu Chen for cryo-EM support; David Houldershaw, Yanni Goudetsidis, and Richard Westlake for computational support; and Gabriel Waksman, Helen R. Saibil and Giulia Zanetti (ISMB) for comments on the manuscript. Cryo-EM data for this investigation were collected at Birkbeck College, University of London with financial support from the Wellcome Trust (202679/Z/16/Z and 206166/Z/17/Z). This work was supported by a Wellcome Trust Senior Research Fellowship to A.J.R. (217186/Z/19/Z) and the BBSRC (BB/S007202/1).

## METHODS

### Construct design for structural studies

Genes encoding the human IFT-A subunits (IFT144/WDR19, IFT140, IFT122, IFT121/WDR35, IFT139/TTC21B, and IFT43) and TULP3 were synthesised with codon optimization for insect cells (Epoch or Twist Bioscience). Genes and expression vectors were amplified using Q5^®^ DNA polymerase (NEB), purified by gel extraction, and combined using HiFi^®^ Assembly Master Mix (NEB). Point mutations were made using the Q5 Site-Directed Mutagenesis Kit (NEB) with mutagenic primers or mutagenic double-stranded DNA fragments (gBlocks^™^, IDT). For insect cell expression, the following constructs were made.

IFT144 was inserted with a C-terminal Tobacco Etch Virus (TEV) protease cleavage site and FLAG tag into the pACEBac1 vector (Geneva Biotech) (IFT144^FLAG^). An IFT144 construct lacking the N-terminal β-propeller domain (residues 2–349 inclusive; IFT144ΔN^FLAG^) was made using the Q5^®^ Site-Directed Mutagenesis Kit (NEB).

IFT140 was inserted with a C-terminal TEV site and Strep-tag^®^II (IFT140^Strep^) or an N-terminal ZZ-tag (^ZZ^IFT140) into the pIDC (Geneva Biotech) or pACEBac1 vector. An IFT140^Strep^ construct with point mutations designed to disrupt the interface with IFT144 (D789R/F792A/K796E /V822A/N826A/A830Y/A833Y/R837E) was made using a gBlock^™^ (IFT140^Mut-Strep^).

A dual expression construct (IFT144^FLAG^-IFT140^Strep^) was generated by fusing the pACEBac1-IFT144^FLAG^ vector and IFT140^Strep^ vector using Cre recombinase according to the MultiBac protocol (Geneva Biotech).

IFT122 was inserted into the pRZ vector (a derivative of pACEBac1; (Toropova et al., 2019)) without tags (IFT122). An IFT122 construct with point mutations designed to disrupt the TULP interface (E1052R/Y1058A/S1062A/E1217R/D1218R) was made using a gBlock^™^ (IFT122^TULPmut^).

IFT139 was inserted into the pACEBac1 vector with an N-terminal His8-ZZ tag, TEV site, and SNAPf^®^ tag (^ZZ-SNAP^IFT139). IFT121 and IFT43 were each inserted into the pACEBac1 vector with an N-terminal His8-ZZ tag and TEV site (^ZZ^IFT121 and ^ZZ^IFT43). For some experiments, an untagged IFT43 construct was used (IFT43). TULP3 was inserted into the pRZ vector with a C-terminal TEV site and His8-ZZ tag (TULP3^ZZ^).

### Protein expression and purification

Protein constructs were expressed in *Spodoptera frugiperda* (Sf9) cells (Gibco) using the baculovirus system as previously described (Toropova et al., 2019), with minor modifications. Virus stocks (V_1_) were generated as described. For co-infections with multiple viruses, small scale (50 ml) cultures were used to optimize relative expression levels. The percentage virus used in each expression is given below. Cultures were incubated for 3 d post infection, then harvested by centrifugation at 2000 g for 30 min at 4 °C. Cell pellets were resuspended in 10 ml 1X PBS (Gibco) per 250 ml culture, then centrifuged at 2000 g for 10 min before flash-freezing in liquid nitrogen. Pellets were either thawed immediately for purification or stored at −80 °C. Purifications were performed at 4 °C. Purified proteins were analyzed by SDS-PAGE on 4–12 % Bis-Tris gels (Thermo Fisher Scientific) with InstantBlue staining (Abcam) and protein identity was verified by mass spectrometry (BSRC Mass Spectrometry and Proteomics Facility, University of St Andrews).

#### Co-purification of IFT144·IFT140·IFT1 22

In a typical preparation, V_1_ baculoviruses for IFT144^FLAG^-IFT140^Strep^ and IFT122 were added to a 2 L Sf9 cell culture at a ratio of 0.25% and 3% [v/v] respectively, incubated for 3 d then harvested and frozen as above. Frozen cell pellets were thawed on ice and resuspended in 80 ml of Buffer A (50 mM Tris [pH 7.5], 150 mM K-acetate, 2 mM Mg-acetate, 1 mM EGTA, 10% [v/v] glycerol, 1 mM DTT, 1 mM PMSF, 0.2 mM Mg-ATP), supplemented with cOmplete^™^ EDTA-free protease inhibitor cocktail (Roche). Resuspended cells were passed 10 times through a pre-cooled Dounce homogeniser with a tight clearance pestle, before clarifying with ultracentrifugation at 183,960 g in a Type 70 Ti rotor for 30 min at 4 °C. The supernatant was passed through a gravity flow column with 1.5 ml bed volume of Strep-Tactin^®^XT resin (IBA) that had been previously equilibrated in Buffer B (50 mM Tris [pH 7.5], 150 mM K-acetate, 2 mM Mg-acetate, 1 mM EGTA, 1 mM DTT, 1 mM PMSF). The protein-bound resin was washed with 10 ml of Buffer B, followed by 5 ml of Buffer B supplemented with 5 mM Mg-ATP. The resin was washed again with 10 ml of Buffer B, and protein was eluted in a series of 1 ml fractions with Buffer B supplemented with 50 mM biotin, with 5 min incubation between each elution. A typical yield was 1 ml of IFT144^FLAG^·IFT140^Strep^·IFT122 subcomplex at a concentration of 0.7 mg/ml (~1.5 μM). Aliquots of 100 μl were flash-frozen in liquid nitrogen and stored at −80 °C. IFT144^FLAG^·IFT140^Strep^·IFT122^TULPmut^ was expressed and purified in the same way.

#### Co-purification of IFTI44ΔNoIFTI4O·IFTI 22

V_1_ baculoviruses for IFT144ΔN^FLAG^, ^ZZ^IFT140 and IFT122 were added to a 500 ml Sf9 cell culture at a ratio of 0.25%, 0.5%, 3% [v/v] respectively, incubated for 3 d then harvested and frozen as above. Frozen cell pellets were thawed on ice and resuspended in 20 ml of IgG Buffer (30 mM HEPES [pH 7.4], 150 mM KCl, 1 mM EGTA, 10% [v/v] glycerol), supplemented with cOmplete™ EDTA-free protease inhibitor cocktail (Roche). Resuspended cells were passed 10 times through a pre-cooled Dounce homogeniser with a tight clearance pestle, before clarifying with ultracentrifugation at 183,960 g in a Type 70 Ti rotor for 30 min at 4 °C. The supernatant was then incubated on a roller for 1 hr with 1 ml IgG Sepharose resin (GE Healthcare) that had been previously equilibrated in IgG Buffer. The resin was then pelleted by centrifugation at 670 g, resuspended in 5 ml of IgG Buffer, and added to a pre-cooled 20 ml gravity flow column (BioRad). The resin was washed twice with 20 ml of IgG Buffer, followed by 25 ml TEV Buffer (50 mM Tris [pH 7.5], 150 mM K-acetate, 2 mM Mg-acetate, 10% [v/v] glycerol, 1 mM EGTA, 1 mM DTT, 1 mM PMSF). Protein-bound resin was resuspended in 1 ml TEV Buffer and transferred to a 2 ml microcentrifuge tube for overnight cleavage with 100 μg of TEV protease. After incubation, the resin was re-applied to a pre-cooled empty column and the cleaved protein was allowed to flow through and collected. Aliquots of 50 μl were flash frozen in liquid nitrogen and stored at −80 °C. A typical yield was 2 ml of IFT144ΔN·IFT140·IFT122 subcomplex at a concentration of 0.25 mg/ml (~0.7 μM). Aliquots of 100 μl were flash-frozen in liquid nitrogen and stored at −80 °C.

#### Co-purification of IFT121·IFT43

V_1_ baculoviruses for ^ZZ^IFT121 and ^ZZ^IFT43 or untagged IFT43 were added to a 2 L Sf9 cell culture at a ratio of 1% and 1.25% [v/v] respectively, incubated for 3 d then harvested and frozen as above. Purifications were performed as for the IFT144ΔNoIFT140·IFT122 subcomplex, with the following alterations. For lysis, cell pellets were lysed in IgG Buffer supplemented with cOmplete^™^ EDTA-free protease inhibitor cocktail (Roche) and 0.2 mM Mg-ATP. The supernatant was incubated with 1 ml of IgG Sepharose resin (GE Healthcare) for 1 hr, before washing twice with 20 ml IgG Buffer, followed by 5 ml of TEV Buffer supplemented with 5 mM Mg-ATP and 25 ml of TEV Buffer, prior to overnight TEV cleavage. IFT121·IFT43 was concentrated to 0.7 mg/ml using an Amicon Ultra centrifugal filter device (100 kDa molecular weight cut-off). Aliquots of 100 μl were flash-frozen in liquid nitrogen and stored at −80 °C. For cryo-EM dataset 1, untagged IFT43 was used. For cryo-EM dataset 2, ^ZZ^IFT43 was used. There was no discernible difference between the datasets.

#### Purification of IFT139

V’baculovirus for ^ZZ-SNAP^IFT139 was added to a 500 ml Sf9 culture at a ratio of 1% [v/v] and incubated for 3 d, then harvested and frozen as above. The cell pellet was thawed, lysed and clarified in IgG Buffer, as described for the IFT144ΔN·IFT140·IFT122. Final IFT139 concentration after TEV cleavage was 2.0 mg/ml. Aliquots of 50 μl were flash-frozen in liquid nitrogen and stored at −80 °C.

#### Purification of TULP3

TULP3^ZZ^ was expressed and purified as for IFT139 except that the starting volume of Sf9 cells was 250 ml. The final concentration of TULP3 after TEV cleavage was 0.6 mg/ml. Aliquots of 50 μl were flash-frozen in liquid nitrogen and stored at −80 °C.

#### IFT144·IFT140 binding assay

V_1_ baculoviruses for IFT144^FLAG^ and IFT140^Strep^/IFT140^Mut-Strep^ were added to a 50 ml Sf9 cell culture at a ratio of 1.25% and 1% [v/v] respectively, incubated for 3 d, then harvested and frozen as above. The cell pellets were thawed in 4 ml Lysis Buffer (50 mM Tris [pH 7.5], 150 mM K-acetate, 2 mM Mg-acetate, 1 mM EGTA, 10% [v/v] glycerol, 1 mM DTT, 1 mM PMSF) supplemented with cOmplete™ EDTA-free protease inhibitor cocktail (Roche). Samples were sonicated (2 s on, 5 s off) for a total of 10 s. Lysates were clarified by centrifugation at 135,557 g in a TLA-110 rotor for 30 min at 4 °C. The supernatant was passed through a gravity flow column with 100 μl bed volume of Strep-Tactin^®^XT resin (IBA), previously equilibrated in Buffer B. The protein-bound resin was washed with 2 ml of Buffer B. Resin was then resuspended in 100 μl Buffer B, and 50 μl was taken and mixed with 16.6 μl NuPAGE^™^ LDS Loading Buffer (Invitrogen). This was heated to 95 °C for 5 min, before taking 5 μl for analysis by SDS-PAGE.

### Reconstitution of the IFT-A complex

Purified IFT144·IFT140·IFT122, IFT121·IFT43, and IFT139 were rapidly thawed and mixed on ice such that the final molar ratio for each protein was approximately one. Following 15 min incubation, the sample was loaded onto a Superose 6 Increase 3.2/300 size exclusion column pre-equilibrated with size-exclusion chromatography (SEC) buffer (50 mM Tris [pH 7.5], 150 mM K-acetate, 2 mM Mg-acetate, 1 mM EGTA, 1 mM DTT) on an ÄKTAmicro system (GE Healthcare). Fractions were analyzed by SDS-PAGE on 4–12 % Bis-Tris gels (Thermo Fisher Scientific) with InstantBlue staining (Abcam). Peak fractions containing all six IFT-A subunits were immediately analyzed by negative stain electron microscopy as described (Toropova et al., 2019) or cryo-EM. The same SEC protocol was used to assay binding of TULP3 to IFT-A and IFT-A(122^TULPmut^) by including purified TULP3 in the reconstitution mix. The same protocol was also used to prepare the IFT-A complex containing IFT144ΔN.

### Cryo-electron microscopy

#### Sample vitrification

Following size-exclusion chromatography, IFT-A complex was vitrified using a Vitrobot Mark IV system (ThermoFisher Scientific) operating at 4 °C and 95–96% humidity. Sample (3 μl) was applied to glow discharged EM grids containing a lacey carbon film covered with a 3 nm layer of continuous carbon (Agar Scientific). Following 30 s of incubation the grid was blotted for 8–12 s using a force setting of −10 before plunging into liquid ethane and storage in liquid nitrogen.

#### Data collection

Three datasets were collected on an in-house Titan Krios instrument (ThermoFisher Scientific) operating at 300 keV and equipped with a BioQuantum K3 direct electron detector and post-column GIF energy filter (slit width; 20 eV) (Gatan, Inc.). Dataset 1 and dataset 2 were of the IFT-A complex containing full-length IFT144. Dataset 3 was of the IFT-A complex containing IFT144ΔN. EPU software operating in “faster acquisition” mode was used for image acquisition. Images were collected at 81,000X nominal magnification as super-resolution, 50-frame movies (0.5335 Å/pixel sampling). Exposure times of 3–3.5 s were used giving electron exposures of 49.5, 48.0 and 47.8 e^-^/Å^2^ for datasets 1–3 respectively. In total 6,888, 10,919 and 7,692 movies were collected for datasets 1–3 respectively using nominal defocus ranges of −1.5 to −3.5 μm.

#### Image pre-processing

Pre-processing was carried out using cryoSPARC v3.2 (Punjani et al., 2017). Movies were aligned, dose-weighted and summed using patch motion-correction. An output F-crop factor of 1/2 was used, which gave a 1.067 Å/pixel sampling for aligned and summed micrographs. CTF parameters were determined using patch CTF estimation with default settings. Dataset 2 micrographs were selected to keep micrographs with a CTF-fit score better than 5 Å. For dataset 3, micrographs with a CTF-fit score better than 5 Å, average intensity of +/-50 and total full-frame motion distance of less than 40 pixels were used.

### IFT-A2 reconstruction

Unless otherwise stated processing was carried out using cryoSPARC v3.2. Blob picker was used on 200 micrographs of dataset 1 using circular blobs of diameter 200–500 Å. Picks were pruned to exclude those with thick carbon background or high intensity regions and extracted into 256 pixel boxes (2.134 Å/pixel sampling) giving a total of 22,826 particles. Following 2D classification, 4 class averages (low-pass filtered to 20 Å) were selected for template-based picking of dataset 1 micrographs using a diameter of 500 Å and particle separation distance of 250 Å. Initial picks (1,438,717) were pruned as described above and 560,406 particles were extracted into 256 pixel boxes (2.134 Å/pixel sampling) followed by 2D classification. Poor quality classes were discarded and a second round of 2D classification performed. From this classification, 7 class averages displaying high detail and different views were selected for *ab initio* 3D structure determination (59,940 particles). This *ab initio* map (low-pass filtered to 30 Å) was used as a starting model for non-uniform 3D refinement of 121,015 particles (2.134 Å/pixel sampling) selected from high-quality 2D classes. The resulting map (low-pass filtered to 30 Å) was used as a starting model for non-uniform 3D refinement of the unbinned particles (1.067 Å/pixel sampling, 512 pixel boxes), yielding a 3.8 Å map.

For dataset 2, three 2D class averages from dataset 1 (low-pass filtered to 20 Å) were used as templates for particle picking. Initial picks (3,707,100) were pruned to 1,121,510 particles as described for dataset 1 and extracted into 256 pixel boxes (2.134 Å/pixel sampling). Two rounds of 2D classification and selection were performed. The dataset 1 map (low-pass filtered to 30 Å) was used a starting model for non-uniform 3D refinement of the unbinned particles (1.067 Å/pixel sampling, 512 pixel boxes), yielding a 3.9 Å map.

The maps from dataset 1 and 2 displayed highly similar features and particles from both datasets (242,645 particles) were combined into a single stack and refined to 3.5 Å resolution using non-uniform 3D refinement (Figure S1D). Local refinements of IFT-A2 were performed using masks displayed in Figure S1H–J using the pose/shift Gaussian prior option. Local resolution estimation was performed using Blocres (cryoSPARC implementation) with default settings.

### IFT-A1 reconstruction

Inspection of 2D classes of the IFT-A complex showed a weak halo of density extending from the IFT-A2 module (Figure S1B), corresponding to the IFT-A1 module that is flexibly attached to IFT-A2. To obtain images centred on IFT-A1 and to prevent IFT-A2 from dominating alignments, we used signal subtraction in RELION v3.1 to remove the IFT-A2 signal. 2D classification on these signal-subtracted particles yielded class averages of the IFT-A1 module in a range of views (Figure S1B), from which 11,490 particles were used to generate an *ab initio* model in cryoSPARC (resolution limit of 15 Å during refinement).

Following identification of the IFT-A1 module, the following strategy was used to maximise the yield of IFT-A1 particles for 3D reconstruction. In cryoSPARC, three IFT-A1 class averages and three IFT-A2 class averages (all low-pass filtered to 20 Å) were used as templates for particle picking. Permissive picking parameters were used (350 Å diameter, 70 Å separation) to generate 7,638,166 initial picks. These were pruned as described above and 1,641,213 particles extracted into 256 pixel boxes (2.134 Å/pixel sampling). These particles were then subjected to supervised 3D classification against maps of IFT-A2 and IFT-A1 (low-pass filtered to 20 Å) using 3D heterogeneous refinement. IFT-A1 particles were isolated by two rounds of 2D classification followed by another round of 3D heterogeneous refinement. Unbinned IFT-A1 particles were extracted into 512 pixel boxes (1.067 Å/pixel sampling) and particles closer than 150 Å were removed. Following a final round of 2D classification, non-uniform 3D refinement on the remaining 68,606 particles was used to give an overall ~10 Å map. The same strategy as described was used for dataset 2 and 68,011 IFT-A1 particles were refined to give another ~10 Å map. IFT-A1 particles from datasets 1 and 2 were then combined (136,617 particles) and refined to give an ~7–15 Å map (Figure S1F). Local resolution estimation was performed using Blocres (cryoSPARC implementation) with default settings.

Essentially the same strategy was used to process the IFT-A1 module in IFT-A(IFT144ΔN) (dataset 3), except IFT-A2 class averages were not used as templates for particle picking. An *ab initio* model was generated from 3 class averages (2,536 particles) and used as a starting model (low-pass filtered to 30 Å) for non-uniform 3D refinement (30,523 particles). This map was then aligned to the IFT-A1 map containing full-length IFT144 using the Fit in Map command in UCSF Chimera (Pettersen et al., 2004) to allow assignment of IFT144 and IFT140 β-propellers.

#### Multi-body analysis

Multi-body analysis of IFT-A cryo-EM data was carried out in RELION v3.1 (Nakane et al., 2018). Dataset 1 IFT-A2 particles (121,015 particles) were imported into RELION and 3D refined against a 30 Å low-pass filtered IFT-A2 reference to give a consensus refinement. These particles were then signal-subtracted to retain the IFT-A1 signal as described. Following 2D classification, particles with clearly resolved IFT-A1 modules were selected (64,792 particles). This identified particles which had clear signal for both IFT-A2 and IFT-A1 modules. These particles were then extracted into larger 350 pixel boxes (2.134 Å/pixel sampling) with no signal subtraction. For multi-body analysis the rlnBodySigmaAngles value was set to 10 and 30; rlnBodySigmaOffset value to 2 and 6 for IFT-A2 and IFT-A1 modules respectively. The output volumes from principle component analysis of the motions were filtered to 30 Å (Movie S1).

### Model building

Atomic model building into the 3.5 Å resolution IFT-A2 map began by using well-resolved side chain densities to build segments of IFT122, IFT121 and IFT139 *de novo* in COOT (Emsley et al., 2010). These models were completed by a combination of manual building in COOT and MDFF of AlphaFold2 (AF) models (Jumper et al., 2021; Varadi et al., 2022) into the cryo-EM map using ISOLDE (Croll, 2018). In general, the local fold of the AF models showed an excellent fit to the experimental density, but the angles of the β-propeller domains and curvature of the long TPR regions required substantial rebuilding, for which we used MDFF with adaptive distance restraints in ISOLDE (Croll and Read, 2021) (Figure S2A). Restraints were then released to inspect and correct outliers, rotamers, and fit-to-density in ISOLDE. For the less well-resolved N-terminal end of IFT139, adaptive distance restraints were retained to prevent over fitting. A path of unfilled density between IFT121 and IFT139 fit the expected position for IFT43. This was independently confirmed by an AF Multimer (Evans et al., 2021) prediction of IFT139·IFT121·IFT43, which exhibited a high confidence score (pLDDT) and low Predicted Aligned Error (PAE) (Figure S2B) and was rebuilt into the density using ISOLDE.

For the 7–15 Å resolution IFT-A1 map, identity of the IFT144 and IFT140 N-terminal β-propellers was established by comparison with the IFT-A1(IFT144ΔN) map (Figure 3C). This assignment was independently confirmed by a high-confidence AF Multimer prediction of the interface between IFT144, IFT140, and C-terminal region of IFT122 (Figure S2C). We docked this model into the IFT-A1 map using a global fitting analysis (Chimera MultiFit) from 10,000 random starting positions, which returned a unique top-scoring fit (cross correlation score 0.83) that agreed with the experimental assignment. AF models for the remaining portions of IFT144 and IFT140 were fit to the density using MDFF with adaptive distance restraints in ISOLDE. To generate the complete model of IFT-A, the IFT-A1 and IFT-A2 models were joined via the overlapping TPR region of IFT122 and flexibly fit to the average IFT-A conformer from RELION multi-body analysis.

To build a pseudo-atomic model of the IFT-A polymer, the IFT-A2 atomic model was docked into the *in situ* sub-tomogram average of the *C. reinhardtii* anterograde IFT-A train (EMDB-4304 (Jordan et al., 2018)) using a global fitting analysis (Chimera MultiFit) from 10,000 random starting positions. This returned a unique top-scoring fit (cross correlation score 0.89) that closely matched the density (Figure 5B). We then placed the complete IFT-A model at these coordinates and used MDFF with adaptive restraints to adjust the IFT-A1 module into the density, yielding an unambiguous fit (Movie S2). The resulting angle between the IFT-A1 and IFT-A2 lies in the range observed in the isolated complex (Figure 1 D–F). A map of the IFT-A polymer was generated by combining repeats of EMDB-4304 (Jordan et al., 2018) using the Chimera “vop maximum” command. Fourteen copies of the IFT-A atomic model were docked into the polymer map using the Chimera “fit” command with the sequential fitting option.

### AlphaFold analysis

AlphaFold2 and AlphaFold Multimer predictions were run using the DeepMind AlphaFold Colab (Evans et al., 2021; Jumper et al., 2021; Varadi et al., 2022) or ColabFold (Mirdita et al., 2022), with Amber relaxation and without the use of templates.

### Visualization

Cryo-EM maps and atomic coordinates were visualized using UCSF Chimera (Pettersen et al., 2004) and UCSF Chimera X (Pettersen et al., 2021).

### Cell culture

IMCD3-FlpIn cells (gift from P.K. Jackson, Stanford University) (Mukhopadhyay et al., 2010) were cultured in DMEM/F-12 media (Gibco) supplemented with 10% Fetal Bovine Serum (FBS) and 100 U/ml penicillin-streptomycin (Gibco). Cells were incubated in serum-free medium for 24 hr to induce ciliogenesis. For Smoothened (Smo) experiments, serum-starved cells were treated with Smo agonist (SAG) (Sigma-Aldrich 566660) at a final concentration of 400 nM for 24 hr. For HaloTag labeling, cell were incubated for 15 min with 1 nM JF646-HaloTag (gift from Luke Lavis; (Grimm et al., 2015)), washed with 1X PBS, then imaged.

### Construct design for cell biology studies

For CRISPR-Cas9 genome editing experiments, the following guide RNAs (gRNAs) targeting IFT144, IFT140, IFT122, IFT121 and IFT139 in IMCD-3 cells were designed using Benchling software and cloned into the pX330 vector (gift from Feng Zhang; Addgene plasmid # 42230; (Cong et al., 2013)): 5’- GAAACTACCTTGCAGTAACA-3’ and 5’- AAAACCAGCCAGCTGGACAA-3’ targeting exons 2 and 4 of IFT144 respectively; gRNAs 5’- TCTATAAGCCCATCTTCTGG-3’ and 5’- TGCCTGACACCCACATTGAG-3’ targeting exons 2 and 3 of IFT140 respectively; 5’-TTCCATCAGGCTTAAATGCG-3’ and 5’-CAAAGACACCGTGTACTGTG-3’ targeting exons 2 and 3 of IFT122 respectively; 5’-TATCCTGGAACAAGGACCAA-3’ and 5’-GTTCATGGAAAGGTTACTAG-3’ targeting exons 2 and 3 of IFT121 respectively; 5’-CACTGGCAACAAGGAGCACG-3’ and 5’-AATGGCCTCAAACTCCCGAA-3’ targeting exons 2 and 3 of IFT139 respectively.

For stable expression of mNeonGreen (mNG) tagged cytoplasmic dynein-2 heavy chain (DYNC2H1) in IMCD3-FlpIn cells, the human DYNC2H1 open reading frame (Toropova et al., 2019) was inserted into the pEF5-FRT vector with N-terminal mNG, FLAG and PreScission Protease (PP) tags.

For stable expression of mNeonGreen (mNG) tagged GPR161 or SSTR3 in IMCD3-FlpIn cells, the GPR161 or SSTR3 open reading frame was inserted into a modified pEF5-FRT vector in which TATA box sequence in the EF1α promoter was mutated to prevent excessive expression that perturbs cilia length (Ye et al., 2018). GPR161 and SSTR3 were tagged with C-terminal TEV, FLAG, PP, and mNG.

For stable expression of mScarlet (mSc) tagged IFT-A proteins (IFT144, IFT140, IFT122, IFT121, and IFT139) and associated mutants using the Super PiggyBac system (System Biosciences), the relevant open reading frames were inserted into the PB-EF1α-MCS-IRES-Neo vector with C-terminal TEV, FLAG, PP, mSc tags, with the exception of IFT139, which we tagged at the N-terminus based on our structure.

For expression of Halo tagged ICK in IMCD3-FlpIn cells, the ICK open reading frame was inserted into a modified pEF5-FRT vector in which TATA box sequence in the EF1α promoter was mutated to prevent excessive expression (Ye et al., 2018) with an N-terminal Halo tag.

### Genome editing and verification

CRISPR-Cas9 genome editing experiments in IMCD-3 cells were performed as described (Perretta-Tejedor et al., 2020) with minor modifications. pX330 vectors expressing the relevant chimeric gRNA and codon-optimized human SpCas9 were transfected into IMCD3-FlpIn cells. In a well of a six well plate, 50–60% confluent cells were transfected with 2.4 μg plasmid DNA using 8.5 μl Lipofectamine^™^ 2000 transfection reagent (Thermo Fisher Scientific) in 300 μl Opti-MEM reduced serum medium (Thermo Fisher Scientific). Total plasmid DNA was made up in equal parts of each expression vector and pAcGFP1-C1 vector (Addgene) (a GFP reporter plasmid) to a total of 2.4 μg (i.e. 1.2 μg each vector for co-transfections). For positive control of transfection, cells were transfected with the pAcGFP1-C1 vector alone (2.4 μg). For negative control, cells were mock-transfected with transfection reagent in Opti-MEM, but no plasmid DNA. Cells successfully transfected, exhibiting GFP fluorescence (GFP+ cells), were selected by fluorescence-activated cell sorting (FACS). A population of GFP+ cells was collected from the group sort and cultured for two weeks before single-cell sorting by FACS. Single-cell-sorted cells were seeded in 96 well plates in 150 μl conditioned medium which was prepared by sterile filtration of a 1:1 mixture of fresh medium, and medium removed from a flask of cells in exponential growth phase (50–70% confluent). Single-cell-sorted colonies were expanded by subculture to increasingly larger culture vessels (24 well plate to 12 well plate to T25 flask). To confirm mutations, genomic DNA was isolated with QuickExtract™ DNA Extraction Solution (Lucigen) and targeted sequences were amplified by PCR. PCR products were cloned into a pJET1.2/blunt vector (ThermoFisher) and sequenced (Eurofins Genomics) to identify clones with insertions or deletions in the IFT-A genes causing premature stop codons (Figure S4).

### Protein expression

To stably express mNG-DYNC2H1, mNG-GPR161 or mNG-SSTR3 in IFT144, IFT140, IFT122, IFT121, or IFT139 KO cell lines, the following protocol was used. Expression plasmid (0.5 μg) for mNG-DYNC2H1, GPR161-mNG or SSTR3-mNG with a hygromycin resistance marker was transfected into the relevant cell line together with 4.5 μg pOG44 (Flp Recombinase expression vector; gift from Peter Jackson) using Lipofectamine_™_ 2000. The medium was supplied with hygromycin (200 μg/ml) 2 days post-transfection and after 2–3 weeks of selection, single colonies were harvested and screened by PCR for successful incorporation of the gene of interest.

To stably express mSc-tagged IFT-A proteins in these cells, we used Super PiggyBac transposon vector system (System Biosciences). Cells in 6-well plates were co-transfected with PiggyBac plasmid containing gene of interest and a geneticin resistance marker (0.5 μg) and 0.2 μg of Super PiggyBac transposase expression vector using Lipofectamine^™^ 2000. Clones were selected using geneticin resistance (500 μg/ml) after 2 d of transfection until confluent, and screened by live-cell TIRF microscopy to confirm expression of both mNG and mSc labeled proteins. For continued culture, growth media containing 500 μg/ml geneticin and 100 μg/ml hygromycin was used.

For expression of Halo-tagged ICK, cells in 35 mm high glass bottom dish were co-transfected with 1 μg of pAcGFP1-C1 reporter plasmid and 1 μg of HaloTag-ICK expression plasmid using Lipofectamine^™^ 2000. Cells were serum starved 24 hr after transfection and imaged by TIRF microscopy after 24 hr of serum starvation.

### TIRF microscopy

All imaging was conducted on an Eclipse Ti-E inverted microscope with a CFI Apo TIRF 1.49 NA oil objective, Perfect Focus System, H-TIRF module, LU-N4 laser unit (Nikon) and a quad band filter set (Chroma). Images were captured on a iXon DU888 Ultra EMCCD camera (Andor), controlled with NIS-Elements AR Software (Nikon). The microscope was kept in a temperature-controlled environmental chamber (Okolab). Files were imported into FIJI (ImageJ, NIH) (Schindelin et al., 2012) for analysis.

### Immunofluorescence

IMCD-3 cells grown on 0.17 mm thick (#1.5) cover glass (VWR) in 12-well plates or 35 mm high glass bottom dishes (Thistle Scientific) were first washed with 1X PBS, followed by two washes in cytoskeletal buffer (10 mM PIPES [pH 6.9], 100 mM NaCl, 300 mM sucrose, 3 mM MgCl_2_) and fixed in 4% paraformaldehyde prepared in cytoskeletal buffer with 0.5% Triton and 5 mM EGTA as described previously (Hua and Ferland, 2017). Next, cells were incubated with blocking solution (3% BSA [w/v] and 2% FBS [v/v] in 1X PBS) for 1 hr at room temperature. Cells were incubated overnight with respective antibodies. Post primary antibody incubation, cells were washed and incubated with corresponding secondary antibody in blocking solution for 1 hr. Cells were stained with DAPI [4,6-diamidino-2-phenylindole] (Fisher Scientific) for nuclear staining and imaged by TIRF microscopy. The following primary antibodies were used at the indicated dilutions: anti-Acetylated tubulin (Sigma Aldrich T6793; 1:2000); anti-gamma tubulin (Sigma Aldrich T6557; 1:500); anti-Smo (Santa Cruz sc-166685; 1:500); anti-ARL13B (Proteintech 17711-1-AP; 1:2000); anti-IFT88 (Proteintech 13967-1-AP; 1:1000). AlexaFluor labeled secondary antibodies (ThermoFisher) were used at 1:500 dilution.

### Live-cell imaging

150,000 cells were seeded onto a 35 mm high glass bottom dish (Thistle Scientific) and grown for 16-24 hr. Subsequently cells were starved for 24 hr in serum-free media to induce ciliogenesis and transferred for live cell imaging by TIRF microscopy with 100 ms exposures at 37 °C. Cells were imaged in cell culture media [DMEM/F12, HEPES with no phenol red] (Gibco) for not more than 1 hr per dish.

### Image analysis

Image analysis was performed using FIJI (ImageJ, NIH) (Schindelin et al., 2012). For measurement of cilia length, cilia were traced using the “segmented line” tool. For analysis of time-averaged fluorescence distributions, time-lapse movies were Z-projected, background subtracted, and cilia were traced using segmented line tool. The “plot profile” tool was used to obtain fluorescence intensities profiles.

### Immunoblotting

Cells were lysed in RIPA buffer (Thermo Scientific) containing 50 mM Tris [pH 8.0], 150 mM sodium chloride, 1.0% Triton X-100, 0.5% sodium deoxycholate, 0.1% SDS. Samples were sonicated (2 s on, 5 s off) for a total of 10 s. Lysates were normalised to a protein concentration of 35 μg by Bradford assay (BioRad). Samples were separated by SDS-PAGE on 4–12% Bis-Tris gels (ThermoFisher Scientific), and wet-transferred to a nitrocellulose membrane (0.45 μM pore size) using fresh transfer buffer (25 mM Tris-base, 192 mM glycine, 10% methanol), and power pack settings of constant 350 mA, 100 V for 45 min. After transfer, membranes were rinsed briefly with TBS (20 mM Tris-base, 150 mM NaCl), before staining with 0.1% Poncaeu S (Sigma) to detect transfer. Membranes were washed 2X for 5 min each using MilliQ water, then for 5 min with TBS-T (20 mM Tris-base, 150 mM NaCl, 0.02% Tween20), before blocking overnight with TBS-T supplemented with 3% [w/v] skimmed milk powder (Milk:TBS-T). Membranes were then washed 2X for 5 min using TBS-T, before incubating with both monoclonal ANTI-FLAG^®^ M2 mouse antibody (1:1000; Sigma-Aldrich) and anti-GAPDH (1:5000; Cell Signaling Technology) in 2% [w/v] Milk:TBS-T for 4 hr at room temperature. After incubation with anti-GAPDH and anti-FLAG primary antibodies, membranes were washed for 4X 15 min with TBS-T, before incubating simultaneously with Goat Anti-Mouse IgG StarBright Blue 700 (1:1000; BioRad) and Goat Anti-Rabbit IgG (H+L) Alexa Fluor 647 (1:1000; Invitrogen) secondary antibodies in 2% [w/v] Milk:TBS-T for 1 hr at room temperature. Blots were then washed with TBS-T for 4X 15 min, before imaging with multiplex settings on a BioRad ChemiDoc MP (filters 715/30 for StarBright Blue 700 detection, and filters 700/50 for Alexa Fluor 647).

### Statistical Analyses

Statistical analyses were performed with GraphPad Prism v8.3.0. Pairwise comparisons were made using Mann-Whitney tests and multiple comparisons were made using one-way ANOVA followed by Kruskal-Wallis tests, which do not assume normal distributions. Superplots were generated as described previously (Lord et al., 2020).

## Data availability

The cryo-EM maps are available from the Electron Microscopy Data Bank under accession codes EMDB-AAAAA (IFT-A2), EMDB-BBBBB (IFT-A1). Coordinates are available from the RCSB Protein Data Bank under accession codes PDB-XXXX (IFT-A2), PDB-YYYY (IFT-A1) and PDB-ZZZZ (IFT-A docked into sub-tomogram average of anterograde IFT train (EMDB-4304; (Jordan et al., 2018)).

## REFERENCES

von Appen, A., Kosinski, J., Sparks, L., Ori, A., DiGuilio, A.L., Vollmer, B., Mackmull, M.-T., Banterle, N., Parca, L., Kastritis, P., et al. (2015). In situ structural analysis of the human nuclear pore complex. Nature 526, 140–143.

Avidor-Reiss, T., Maer, A.M., Koundakjian, E., Polyanovsky, A., Keil, T., Subramaniam, S., and Zuker, C.S. (2004). Decoding cilia function: defining specialized genes required for compartmentalized cilia biogenesis. Cell 117, 527–539.

Badgandi, H.B., Hwang, S.-H., Shimada, I.S., Loriot, E., and Mukhopadhyay, S. (2017). Tubby family proteins are adapters for ciliary trafficking of integral membrane proteins. J. Cell Biol. 216, 743–760.

Barbeito, P., and Garcia-Gonzalo, F.R. (2021). HTR6 and SSTR3 targeting to primary cilia. Biochem. Soc. Trans. 49, 79–91.

Beck, M., Mosalaganti, S., and Kosinski, J. (2018). From the resolution revolution to evolution: structural insights into the evolutionary relationships between vesicle coats and the nuclear pore. Curr. Opin. Struct. Biol. 52, 32–40.

Behal, R.H., Miller, M.S., Qin, H., Lucker, B.F., Jones, A., and Cole, D.G. (2012). Subunit interactions and organization of the Chlamydomonas reinhardtii intraflagellar transport complex A proteins. J. Biol. Chem. 287, 11689–11703.

Bhogaraju, S., Taschner, M., Morawetz, M., Basquin, C., and Lorentzen, E. (2011). Crystal structure of the intraflagellar transport complex 25/27. EMBO J. 30, 1907–1918.

Bhogaraju, S., Cajanek, L., Fort, C., Blisnick, T., Weber, K., Taschner, M., Mizuno, N., Lamla, S., Bastin, P., Nigg, E.A., et al. (2013). Molecular basis of tubulin transport within the cilium by IFT74 and IFT81. Science 341, 1009–1012.

Bredrup, C., Saunier, S., Oud, M.M., Fiskerstrand, T., Hoischen, A., Brackman, D., Leh, S.M., Midtbø, M., Filhol, E., Bole-Feysot, C., et al. (2011). Ciliopathies with skeletal anomalies and renal insufficiency due to mutations in the IFT-A gene WDR19. Am. J. Hum. Genet. 89, 634–643.

Broekhuis, J.R., Verhey, K.J., and Jansen, G. (2014). Regulation of cilium length and intraflagellar transport by the RCK-kinases ICK and MOK in renal epithelial cells. PLoS One 9, e108470.

Chaya, T., Omori, Y., Kuwahara, R., and Furukawa, T. (2014). ICK is essential for cell typespecific ciliogenesis and the regulation of ciliary transport. EMBO J. 33, 1227–1242.

Chien, A., Shih, S.M., Bower, R., Tritschler, D., Porter, M.E., and Yildiz, A. (2017). Dynamics of the IFT machinery at the ciliary tip. Elife 6, e28606.

Cole, D.G., Diener, D.R., Himelblau, A.L., Beech, P.L., Fuster, J.C., and Rosenbaum, J.L. (1998). Chlamydomonas kinesin-II-dependent intraflagellar transport (IFT): IFT particles contain proteins required for ciliary assembly in Caenorhabditis elegans sensory neurons. J. Cell Biol. 141, 993–1008.

Cong, L., Ran, F.A., Cox, D., Lin, S., Barretto, R., Habib, N., Hsu, P.D., Wu, X., Jiang, W., Marraffini, L.A., et al. (2013). Multiplex genome engineering using CRISPR/Cas systems. Science 339, 819–823.

Croll, T.I. (2018). ISOLDE: a physically realistic environment for model building into low-resolution electron-density maps. Acta Crystallogr. D Struct. Biol. 74, 519–530.

Croll, T.I., and Read, R.J. (2021). Adaptive Cartesian and torsional restraints for interactive model rebuilding. Acta Crystallogr. D Struct. Biol. 77, 438–446.

van Dam, T.J.P., Townsend, M.J., Turk, M., Schlessinger, A., Sali, A., Field, M.C., and Huynen, M.A. (2013). Evolution of modular intraflagellar transport from a coatomer-like progenitor. Proc. Natl. Acad. Sci. U. S. A. 110, 6943–6948.

Devos, D., Dokudovskaya, S., Alber, F., Williams, R., Chait, B.T., Sali, A., and Rout, M.P. (2004). Components of coated vesicles and nuclear pore complexes share a common molecular architecture. PLoS Biol. 2, e380.

Dewees, S.I., Vargová, R., Hardin, K.R., Turn, R.E., Devi, S., Linnert, J., Wolfrum, U., Caspary, T., Eliáš, M., and Kahn, R.A. (2022). Phylogenetic profiling and cellular analyses of ARL16 reveal roles in traffic of IFT140 and INPP5E. Mol. Biol. Cell 33, ar33.

Dodonova, S.O., Diestelkoetter-Bachert, P., von Appen, A., Hagen, W.J.H., Beck, R., Beck, M., Wieland, F., and Briggs, J.A.G. (2015). VESICULAR TRANSPORT. A structure of the COPI coat and the role of coat proteins in membrane vesicle assembly. Science 349, 195–198.

Dodonova, S.O., Aderhold, P., Kopp, J., Ganeva, I., Röhling, S., Hagen, W.J.H., Sinning, I., Wieland, F., and Briggs, J.A.G. (2017). 9Å structure of the COPI coat reveals that the Arf1 GTPase occupies two contrasting molecular environments. Elife 6, e26691.

Emsley, P., Lohkamp, B., Scott, W.G., and Cowtan, K. (2010). Features and development of coot. Acta Crystallogr. D Biol. Crystallogr. 66, 486–501.

Evans, R., O’Neill, M., Pritzel, A., Antropova, N., Senior, A., Green, T., Žídek, A., Bates, R., Blackwell, S., Yim, J., et al. (2021). Protein complex prediction with AlphaFold-Multimer. bioRxiv, https://doi.org/10.1101/2021.10.04.463034

Fotin, A., Cheng, Y., Sliz, P., Grigorieff, N., Harrison, S.C., Kirchhausen, T., and Walz, T. (2004). Molecular model for a complete clathrin lattice from electron cryomicroscopy. Nature 432, 573–579.

Fu, W., Wang, L., Kim, S., Li, J., and Dynlacht, B.D. (2016). Role for the IFT-A complex in selective transport to the primary cilium. Cell Rep. 17, 1505–1517.

Garcia-Gonzalo, F.R., and Reiter, J.F. (2017). Open sesame: How transition fibers and the transition zone control ciliary composition. Cold Spring Harb. Perspect. Biol. 9, a028134.

Grimm, J.B., English, B.P., Chen, J., Slaughter, J.P., Zhang, Z., Revyakin, A., Patel, R., Macklin, J.J., Normanno, D., Singer, R.H., et al. (2015). A general method to improve fluorophores for live-cell and single-molecule microscopy. Nat. Methods 12, 244–250.

Han, S., Miyoshi, K., Shikada, S., Amano, G., Wang, Y., Yoshimura, T., and Katayama, T. (2019). TULP3 is required for localization of membrane-associated proteins ARL13B and INPP5E to primary cilia. Biochem. Biophys. Res. Commun. 509, 227–234.

Hirano, T., Katoh, Y., and Nakayama, K. (2017). Intraflagellar transport-A complex mediates ciliary entry and retrograde trafficking of ciliary G protein-coupled receptors. Mol. Biol. Cell 28, 429–439.

van den Hoek, H., Klena, N., Jordan, M.A., Alvarez Viar, G., Righetto, R.D., Schaffer, M., Erdmann, P.S., Wan, W., Geimer, S., Plitzko, J.M., et al. (2022). In situ architecture of the ciliary base reveals the stepwise assembly of intraflagellar transport trains. Science 377, 543–548.

Hua, K., and Ferland, R.J. (2017). Fixation methods can differentially affect ciliary protein immunolabeling. Cilia 6, 5.

Hunter, M.R., Scourfield, E.J., Emmott, E., and Graham, S.C. (2017). VPS18 recruits VPS41 to the human HOPS complex via a RING-RING interaction. Biochem. J. 474, 3615–3626.

Iomini, C., Babaev-Khaimov, V., Sassaroli, M., and Piperno, G. (2001). Protein particles in Chlamydomonas flagella undergo a transport cycle consisting of four phases. J. Cell Biol. 153, 13–24.

Ishida, Y., Kobayashi, T., Chiba, S., Katoh, Y., and Nakayama, K. (2021). Molecular basis of ciliary defects caused by compound heterozygous IFT144/WDR19 mutations found in cranioectodermal dysplasia. Hum. Mol. Genet. 30, 213–225.

Jackson, L.P., Lewis, M., Kent, H.M., Edeling, M.A., Evans, P.R., Duden, R., and Owen, D.J. (2012). Molecular basis for recognition of dilysine trafficking motifs by COPI. Dev. Cell 23, 1255–1262.

Jakobi, A.J., Wilmanns, M., and Sachse, C. (2017). Model-based local density sharpening of cryo-EM maps. Elife 6, e27131.

Jékely, G., and Arendt, D. (2006). Evolution of intraflagellar transport from coated vesicles and autogenous origin of the eukaryotic cilium. Bioessays 28, 191–198.

Jordan, M.A., and Pigino, G. (2021). The structural basis of intraflagellar transport at a glance. J. Cell Sci. 134, jcs247163.

Jordan, M.A., Diener, D.R., Stepanek, L., and Pigino, G. (2018). The cryo-EM structure of intraflagellar transport trains reveals how dynein is inactivated to ensure unidirectional anterograde movement in cilia. Nat. Cell Biol. 20, 1250–1255.

Jumper, J., Evans, R., Pritzel, A., Green, T., Figurnov, M., Ronneberger, O., Tunyasuvunakool, K., Bates, R., Žídek, A., Potapenko, A., et al. (2021). Highly accurate protein structure prediction with AlphaFold. Nature 596, 583–589.

Katoh, Y., Terada, M., Nishijima, Y., Takei, R., Nozaki, S., Hamada, H., and Nakayama, K. (2016). Overall architecture of the intraflagellar transport (IFT)-B complex containing Cluap1/IFT38 as an essential component of the IFT-B peripheral subcomplex. J. Biol. Chem. 291, 10962–10975.

Kaur, G., and Subramanian, S. (2015). A novel RING finger in the C-terminal domain of the coatomer protein α-COP. Biol. Direct 10, 70.

Kobayashi, T., Ishida, Y., Hirano, T., Katoh, Y., and Nakayama, K. (2021). Cooperation of the IFT-A complex with the IFT-B complex is required for ciliary retrograde protein trafficking and GPCR import. Mol. Biol. Cell 32, 45–56.

Kozminski, K.G., Johnson, K.A., Forscher, P., and Rosenbaum, J.L. (1993). A motility in the eukaryotic flagellum unrelated to flagellar beating. Proc. Natl. Acad. Sci. U. S. A. 90, 5519–5523.

Kozminski, K.G., Beech, P.L., and Rosenbaum, J.L. (1995). The Chlamydomonas kinesin-like protein FLA10 is involved in motility associated with the flagellar membrane. J. Cell Biol. 131, 1517–1527.

Lacey, S.E., Foster, H.E., and Pigino, G. (2022). The Molecular Structure of Anterograde Intraflagellar transport trains. bioRxiv, https://doi.org/10.1101/2022.08.01.502329.

Langousis, G., Cavadini, S., Boegholm, N., Lorentzen, E., Kempf, G., and Matthias, P. (2022). Structure of the ciliogenesis-associated CPLANE complex. Sci. Adv. 8, eabn0832.

Lechtreck, K.F. (2015). IFT-cargo interactions and protein transport in cilia. Trends Biochem. Sci. 40, 765–778.

Lee, C., and Goldberg, J. (2010). Structure of coatomer cage proteins and the relationship among COPI, COPII, and clathrin vesicle coats. Cell 142, 123–132.

Liang, Y., Pang, Y., Wu, Q., Hu, Z., Han, X., Xu, Y., Deng, H., and Pan, J. (2014). FLA8/KIF3B phosphorylation regulates kinesin-II interaction with IFT-B to control IFT entry and turnaround. Dev. Cell 30, 585–597.

Liem, K.F., Jr, Ashe, A., He, M., Satir, P., Moran, J., Beier, D., Wicking, C., and Anderson, K.V. (2012). The IFT-A complex regulates Shh signaling through cilia structure and membrane protein trafficking. J. Cell Biol. 197, 789–800.

Lord, S.J., Velle, K.B., Mullins, R.D., and Fritz-Laylin, L.K. (2020). SuperPlots: Communicating reproducibility and variability in cell biology. J. Cell Biol. 219, e202001064.

Ma, W., and Goldberg, J. (2013). Rules for the recognition of dilysine retrieval motifs by coatomer. EMBO J. 32, 926–937.

McCafferty, C.L., Papoulas, O., Jordan, M.A., Hoogerbrugge, G., Nichols, C., Pigino, G., Taylor, D.W., Wallingford, J.B., and Marcotte, E.M. (2022). Integrative modeling reveals the molecular architecture of the Intraflagellar Transport A (IFT-A) complex. bioRxiv, https://doi.org/10.1101/2022.07.05.498886.

Mirdita, M., Schütze, K., Moriwaki, Y., Heo, L., Ovchinnikov, S., and Steinegger, M. (2022). ColabFold: making protein folding accessible to all. Nat. Methods 19, 679–682.

Mitchison, H.M., and Valente, E.M. (2017). Motile and non-motile cilia in human pathology: from function to phenotypes. J. Pathol. 241, 294–309.

Mukhopadhyay, S., and Jackson, P.K. (2011). The tubby family proteins. Genome Biol. 12, 225.

Mukhopadhyay, S., Wen, X., Chih, B., Nelson, C.D., Lane, W.S., Scales, S.J., and Jackson, P.K. (2010). TULP3 bridges the IFT-A complex and membrane phosphoinositides to promote trafficking of G protein-coupled receptors into primary cilia. Genes Dev. 24, 2180–2193.

Mukhopadhyay, S., Wen, X., Ratti, N., Loktev, A., Rangell, L., Scales, S.J., and Jackson, P.K. (2013). The ciliary G-protein-coupled receptor Gpr161 negatively regulates the Sonic hedgehog pathway via cAMP signaling. Cell 152, 210–223.

Nachury, M.V. (2018). The molecular machines that traffic signaling receptors into and out of cilia. Curr. Opin. Cell Biol. 51, 124–131.

Nachury, M.V., and Mick, D.U. (2019). Establishing and regulating the composition of cilia for signal transduction. Nat. Rev. Mol. Cell Biol. 20, 389–405.

Nachury, M.V., Seeley, E.S., and Jin, H. (2010). Trafficking to the ciliary membrane: how to get across the periciliary diffusion barrier? Annu. Rev. Cell Dev. Biol. 26, 59–87.

Nakamura, K., Noguchi, T., Takahara, M., Omori, Y., Furukawa, T., Katoh, Y., and Nakayama, K. (2020). Anterograde trafficking of ciliary MAP kinase-like ICK/CILK1 by the intraflagellar transport machinery is required for intraciliary retrograde protein trafficking. J. Biol. Chem. 295, 13363–13376.

Nakane, T., Kimanius, D., Lindahl, E., and Scheres, S.H. (2018). Characterisation of molecular motions in cryo-EM single-particle data by multi-body refinement in RELION. Elife 7, e36861.

Nievergelt, A.P., Zykov, I., Diener, D., Chhatre, A., Buchholz, T.-O., Delling, M., Diez, S., Jug, F., Štěpánek, L., and Pigino, G. (2022). Conversion of anterograde into retrograde trains is an intrinsic property of intraflagellar transport. Curr. Biol. S0960-9822(22)01160-5.

Oh, Y.S., Wang, E.J., Gailey, C.D., Brautigan, D.L., Allen, B.L., and Fu, Z. (2019). Ciliopathy-associated protein kinase ICK requires its non-catalytic carboxyl-terminal domain for regulation of ciliogenesis. Cells 8, 677.

Paige Taylor, S., Kunova Bosakova, M., Varecha, M., Balek, L., Barta, T., Trantirek, L., Jelinkova, I., Duran, I., Vesela, I., Forlenza, K.N., et al. (2016). An inactivating mutation in intestinal cell kinase, ICK, impairs hedgehog signalling and causes short rib-polydactyly syndrome. Hum. Mol. Genet. 25, 3998–4011.

Palicharla, V.R., Hwang, S.-H., Somatilaka, B.N., Badgandi, H.B., Legué, E., Tran, V.M., Woodruff, J.B., Liem, K.F., Jr, and Mukhopadhyay, S. (2021). Interactions between TULP3 tubby domain cargo site and ARL13B amphipathic helix promote lipidated protein transport to cilia. bioRxiv, https://doi.org/10.1101/2021.05.25.445488.

Pazour, G.J., Dickert, B.L., and Witman, G.B. (1999). The DHC1b (DHC2) isoform of cytoplasmic dynein is required for flagellar assembly. J. Cell Biol. 144, 473–481.

Pedersen, L.B., Geimer, S., and Rosenbaum, J.L. (2006). Dissecting the molecular mechanisms of intraflagellar transport in chlamydomonas. Curr. Biol. 16, 450–459.

Perretta-Tejedor, N., Freke, G., Seda, M., Long, D.A., and Jenkins, D. (2020). Generating mutant renal cell lines using CRISPR technologies. Methods Mol. Biol. 2067, 323–340.

Pettersen, E.F., Goddard, T.D., Huang, C.C., Couch, G.S., Greenblatt, D.M., Meng, E.C., and Ferrin, T.E. (2004). UCSF Chimera--a visualization system for exploratory research and analysis. J. Comput. Chem. 25, 1605–1612.

Pettersen, E.F., Goddard, T.D., Huang, C.C., Meng, E.C., Couch, G.S., Croll, T.I., Morris, J.H., and Ferrin, T.E. (2021). UCSF ChimeraX: Structure visualization for researchers, educators, and developers. Protein Sci. 30, 70–82.

Picariello, T., Brown, J.M., Hou, Y., Swank, G., Cochran, D.A., King, O.D., Lechtreck, K., Pazour, G.J., and Witman, G.B. (2019). A global analysis of IFT-A function reveals specialization for transport of membrane-associated proteins into cilia. J. Cell Sci. 132, jcs.220749.

Pigino, G., Geimer, S., Lanzavecchia, S., Paccagnini, E., Cantele, F., Diener, D.R., Rosenbaum, J.L., and Lupetti, P. (2009). Electron-tomographic analysis of intraflagellar transport particle trains in situ. J. Cell Biol. 187, 135–148.

Piperno, G., and Mead, K. (1997). Transport of a novel complex in the cytoplasmic matrix of Chlamydomonas flagella. Proc. Natl. Acad. Sci. U. S. A. 94, 4457–4462.

Piperno, G., Siuda, E., Henderson, S., Segil, M., Vaananen, H., and Sassaroli, M. (1998). Distinct mutants of retrograde intraflagellar transport (IFT) share similar morphological and molecular defects. J. Cell Biol. 143, 1591–1601.

Porter, M.E., Bower, R., Knott, J.A., Byrd, P., and Dentler, W. (1999). Cytoplasmic dynein heavy chain 1b is required for flagellar assembly in Chlamydomonas. Mol. Biol. Cell 10, 693–712.

Punjani, A., Rubinstein, J.L., Fleet, D.J., and Brubaker, M.A. (2017). cryoSPARC: algorithms for rapid unsupervised cryo-EM structure determination. Nat. Methods 14, 290–296.

Quidwai, T., Wang, J., Hall, E.A., Petriman, N.A., Leng, W., Kiesel, P., Wells, J.N., Murphy, L.C., Keighren, M.A., Marsh, J.A., et al. (2021). A WDR35-dependent coat protein complex transports ciliary membrane cargo vesicles to cilia. Elife 10, e69786.

Reiter, J.F., and Leroux, M.R. (2017). Genes and molecular pathways underpinning ciliopathies. Nat. Rev. Mol. Cell Biol. 18, 533–547.

Rosenbaum, J.L., and Witman, G.B. (2002). Intraflagellar transport. Nat. Rev. Mol. Cell Biol. 3, 813–825.

Santagata, S., Boggon, T.J., Baird, C.L., Gomez, C.A., Zhao, J., Shan, W.S., Myszka, D.G., and Shapiro, L. (2001). G-protein signaling through tubby proteins. Science 292, 2041–2050.

Scheidel, N., and Blacque, O.E. (2018). Intraflagellar transport complex A genes differentially regulate cilium formation and transition zone gating. Curr. Biol. 28, 3279–3287.e2.

Schindelin, J., Arganda-Carreras, I., Frise, E., Kaynig, V., Longair, M., Pietzsch, T., Preibisch, S., Rueden, C., Saalfeld, S., Schmid, B., et al. (2012). Fiji: an open-source platform for biological-image analysis. Nat. Methods 9, 676–682.

Schwartz, T.U. (2022). Solving the nuclear pore puzzle. Science 376, 1158–1159.

Sigg, M.A., Menchen, T., Lee, C., Johnson, J., Jungnickel, M.K., Choksi, S.P., Garcia, G., 3rd, Busengdal, H., Dougherty, G.W., Pennekamp, P., et al. (2017). Evolutionary proteomics uncovers ancient associations of cilia with signaling pathways. Dev. Cell 43, 744–762.e11.

Signor, D., Wedaman, K.P., Orozco, J.T., Dwyer, N.D., Bargmann, C.I., Rose, L.S., and Scholey, J.M. (1999). Role of a class DHC1b dynein in retrograde transport of IFT motors and IFT raft particles along cilia, but not dendrites, in chemosensory neurons of living Caenorhabditis elegans. J. Cell Biol. 147, 519–530.

Stepanek, L., and Pigino, G. (2016). Microtubule doublets are double-track railways for intraflagellar transport trains. Science 352, 721–724.

Takahara, M., Katoh, Y., Nakamura, K., Hirano, T., Sugawa, M., Tsurumi, Y., and Nakayama, K. (2018). Ciliopathy-associated mutations of IFT122 impair ciliary protein trafficking but not ciliogenesis. Hum. Mol. Genet. 27, 516–528.

Taschner, M., and Lorentzen, E. (2016). The intraflagellar transport machinery. Cold Spring Harb. Perspect. Biol. 8, a028092.

Taschner, M., Bhogaraju, S., and Lorentzen, E. (2012). Architecture and function of IFT complex proteins in ciliogenesis. Differentiation 83, S12–22.

Taschner, M., Kotsis, F., Braeuer, P., Kuehn, E.W., and Lorentzen, E. (2014). Crystal structures of IFT70/52 and IFT52/46 provide insight into intraflagellar transport B core complex assembly. J. Cell Biol. 207, 269–282.

Taschner, M., Weber, K., Mourão, A., Vetter, M., Awasthi, M., Stiegler, M., Bhogaraju, S., and Lorentzen, E. (2016). Intraflagellar transport proteins 172, 80, 57, 54, 38, and 20 form a stable tubulin-binding IFT-B2 complex. EMBO J. 35, 773–790.

Taschner, M., Lorentzen, A., Mourão, A., Collins, T., Freke, G.M., Moulding, D., Basquin, J., Jenkins, D., and Lorentzen, E. (2018). Crystal structure of intraflagellar transport protein 80 reveals a homo-dimer required for ciliogenesis. Elife 7, e33067.

Toriyama, M., University of Washington Center for Mendelian Genomics, Lee, C., Taylor, S. P., Duran, I., Cohn, D.H., Bruel, A.-L., Tabler, J.M., Drew, K., Kelly, M.R., et al. (2016). The ciliopathy-associated CPLANE proteins direct basal body recruitment of intraflagellar transport machinery. Nat. Genet. 48, 648–656.

Toropova, K., Zalyte, R., Mukhopadhyay, A.G., Mladenov, M., Carter, A.P., and Roberts, A.J. (2019). Structure of the dynein-2 complex and its assembly with intraflagellar transport trains. Nat. Struct. Mol. Biol. 26, 823–829.

Valenstein, M.L., Rogala, K.B., Lalgudi, P.V., Brignole, E.J., Gu, X., Saxton, R.A., Chantranupong, L., Kolibius, J., Quast, J.-P., and Sabatini, D.M. (2022). Structure of the nutrient-sensing hub GATOR2. Nature 607, 610–616.

Varadi, M., Anyango, S., Deshpande, M., Nair, S., Natassia, C., Yordanova, G., Yuan, D., Stroe, O., Wood, G., Laydon, A., et al. (2022). AlphaFold Protein Structure Database: massively expanding the structural coverage of protein-sequence space with high-accuracy models. Nucleic Acids Res. 50, D439–D444.

Wachter, S., Jung, J., Shafiq, S., Basquin, J., Fort, C., Bastin, P., and Lorentzen, E. (2019). Binding of IFT22 to the intraflagellar transport complex is essential for flagellum assembly. EMBO J. 38, e101251.

Webb, S., Mukhopadhyay, A.G., and Roberts, A.J. (2020). Intraflagellar transport trains and motors: Insights from structure. Semin. Cell Dev. Biol. 107, 82–90.

Williamson, S.M., Silva, D.A., Richey, E., and Qin, H. (2012). Probing the role of IFT particle complex A and B in flagellar entry and exit of IFT-dynein in Chlamydomonas. Protoplasma 249, 851–856.

Wingfield, J.L., Mengoni, I., Bomberger, H., Jiang, Y.-Y., Walsh, J.D., Brown, J.M., Picariello, T., Cochran, D.A., Zhu, B., Pan, J., et al. (2017). IFT trains in different stages of assembly queue at the ciliary base for consecutive release into the cilium. Elife 6, e26609.

Wingfield, J.L., Mekonnen, B., Mengoni, I., Liu, P., Jordan, M., Diener, D., Pigino, G., and Lechtreck, K. (2021). In vivo imaging shows continued association of several IFT-A, IFT-B and dynein complexes while IFT trains U-turn at the tip. J. Cell Sci. 134, e259010.

Ye, F., Nager, A.R., and Nachury, M.V. (2018). BBSome trains remove activated GPCRs from cilia by enabling passage through the transition zone. J. Cell Biol. 217, 1847–1868.

Yi, P., Li, W.-J., Dong, M.-Q., and Ou, G. (2017). Dynein-driven retrograde intraflagellar transport is triphasic in C. elegans sensory cilia. Curr. Biol. 27, 1448–1461.e7.

Zhang, Z., Danné, N., Meddens, B., Heller, I., and Peterman, E.J.G. (2021). Direct imaging of intraflagellar-transport turnarounds reveals that motors detach, diffuse, and reattach to opposite-direction trains. Proc. Natl. Acad. Sci. U. S. A. 118, e2115089118.

Zhu, B., Zhu, X., Wang, L., Liang, Y., Feng, Q., and Pan, J. (2017). Functional exploration of the IFT-A complex in intraflagellar transport and ciliogenesis. PLoS Genet. 13, e1006627.

