## Supplemental Information for "IFT-A Structure Reveals Carriages for Membrane Protein Transport into Cilia"

[#]These authors contributed equally

**SUPPLEMENTAL INFORMATION**

Figure S1–S6

Table S1–S3

Movie S1 & S2 legends

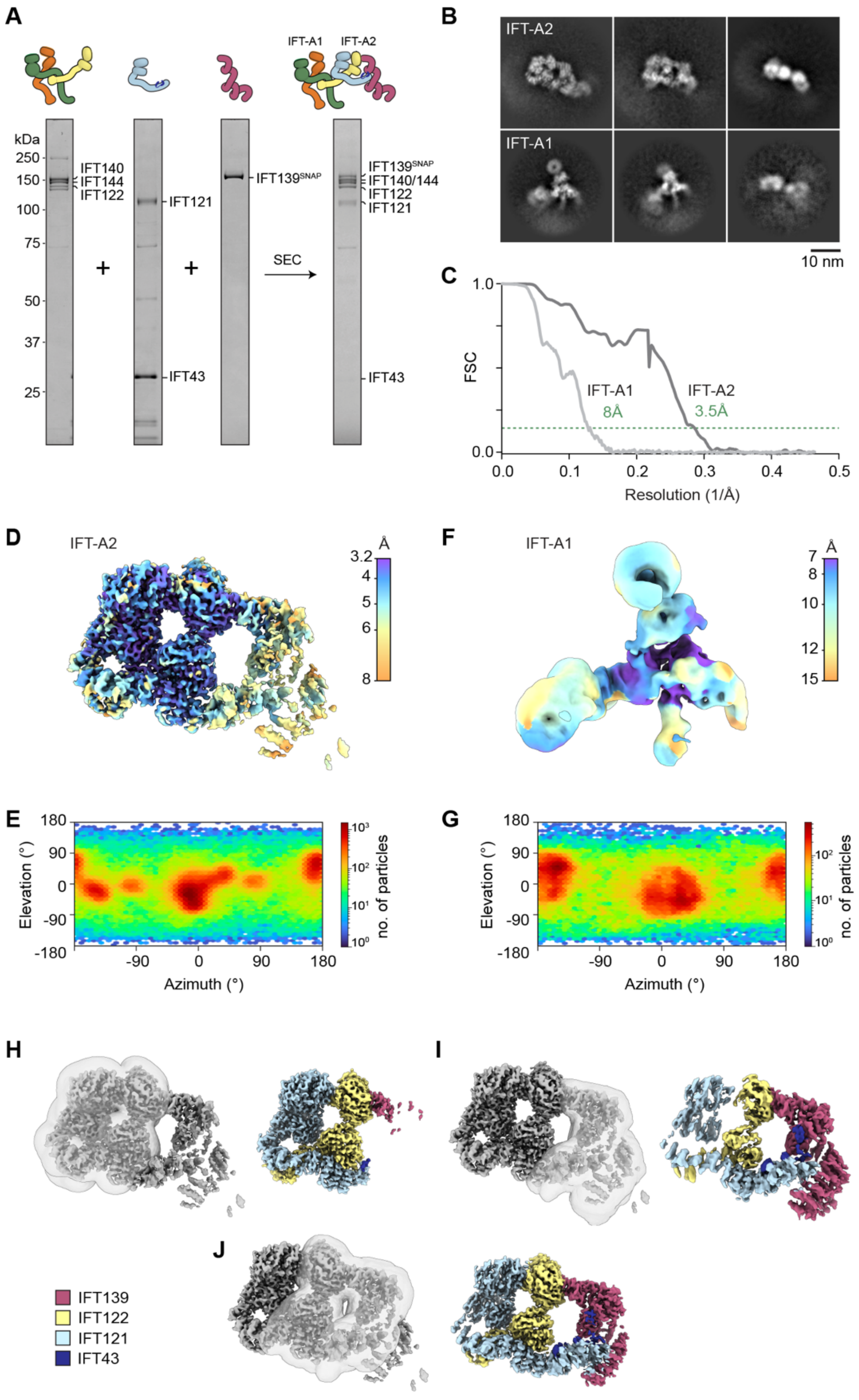


**Figure S1 – Reconstitution and Cryo-EM of the Human IFT-A Complex**

(A) SDS-PAGE of purified human IFT-A components (left three lanes) used to reconstitute the IFT-A complex (right lane). Complex was isolated using size-exclusion chromatography (SEC).

(B) Cryo-EM class averages of IFT-A2 (top row) and IFT-A1 (bottom row).

(C) Fourier shell correlation (FSC) plots for the IFT-A2 and IFT-A1 reconstructions. Resolution at FSC=0.143 is marked.

(D) IFT-A2 reconstruction colored by local resolution, calculated using Blocres (cryoSPARC implementation). Map sharpened with global B-factor of -102 Å$^2$.

(E) Angular distribution of IFT-A2 particles.

(F) IFT-A1 reconstruction colored by local resolution, calculated as for (D). Unsharpened map shown.

(G) Angular distribution of IFT-A1 particles.

(H–J) Local 3D refinement of different regions in the IFT-A2 module. For each region, the mask applied to the reference during refinement is shown on the left, with the refined and sharpened map colored by subunit on the right. Global B-factors of -58, -97 and -60 Å$^2$ were applied for (H), (I) and (J) respectively.

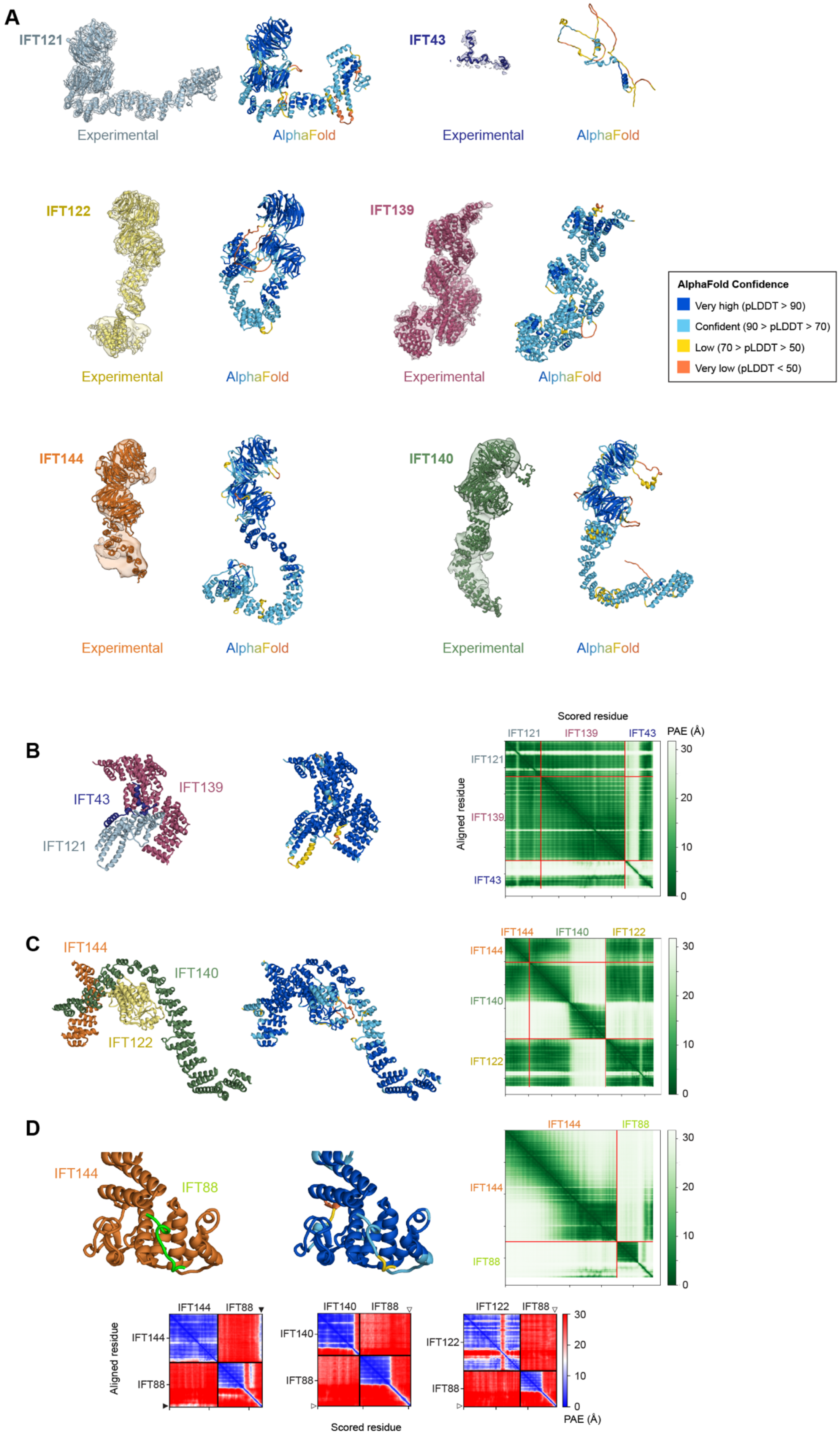


**Figure S2 – AlphaFold Analysis of IFT-A and -B** (legend continued overleaf)

**Figure S2 – AlphaFold Analysis of IFT-A and -B** (legend continued)
(A) Comparison of the IFT-A subunit structures and conformations obtained in our structure (colored by subunit, cryo-EM density shown) with raw AlphaFold (AF) models. AF models colored by pLDDT (predicted local distance difference test) according to the key.
(B) AF Multimer prediction of the interface between IFT139, IFT121, and IFT43 in ribbon representation. Left, colored by subunit; right, by pLDDT confidence. Far right, predicted aligned error (PAE) plot. Note the high confidence prediction (low PAE) for the interface between the C-terminal region of IFT43 and IFT139 and IFT43. The unstructured N-terminal region of IFT43 not shown for clarity.
(C) AF Multimer prediction of the interface between IFT144, IFT140 and IFT122. Coloring and PAE plot as for (B). Note the high confidence prediction (low PAE) for interface between the IFT122 CTD and IFT144 and IFT140 TPR regions.
(D) AF Multimer prediction of the interface between IFT144 and C-terminal loop of IFT88 (rest of IFT88 not shown). Note the high confidence prediction (low PAE) for the small interface between the IFT88 C-terminal loop IFT144 TPR region. To establish the specificity of this small interface, we formed control AF predictions between IFT88 and the TPR regions of IFT144, IFT140 and IFT122 (bottom row). These predictions were performed using ColabFold with MM2Seqs rather than the DeepMind AlphaFold Colab for comparison (hence different coloring of the PAE plots). Interaction is predicted between the IFT88 C-terminal loop and IFT144 (black arrowheads, low PAE), but not with IFT140 or IFT122 (white arrowheads, high PAE).

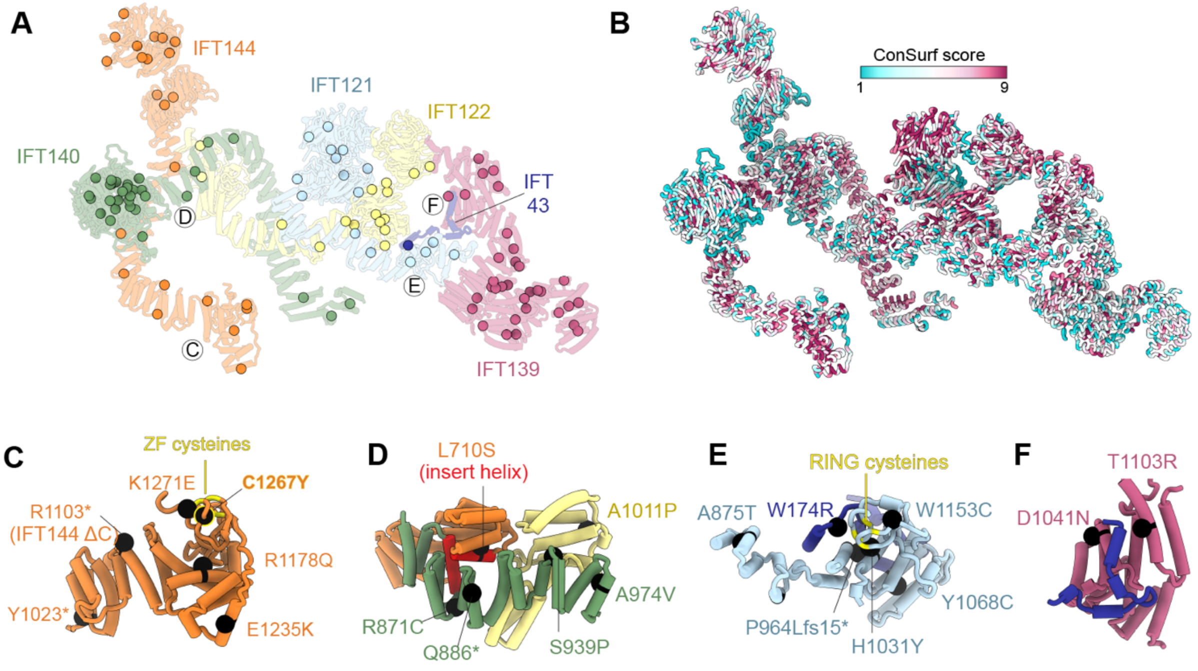


**Figure S3 – Analysis of IFT-A Structure and Disease Mutations**
(A) Curated list of IFT-A missense, nonsense and frameshift mutations (see Table S3) plotted onto the structure of IFT-A. Spheres depict the Cβ atom of the mutated residue, colored by subunit. For frameshift mutations, the first altered residue is depicted. Many disease mutations map to internal sites within β-propeller domains and TPR repeats and are likely to cause folding defects. Others map to subunit interfaces and may thus perturb IFT-A complex formation or function (see C–F).
(B) IFT-A structure in ribbon representation colored by sequence conservation (1– low; 9 – high), calculated using the ConSurf server (Landau et al., Nucleic Acids Res. 33:W299-302, 2005).
(C–F) Close up views of selected mutations occurring at subunit interfaces and domains labeled in A (circled letters) and mentioned in the main text. R1103* mutation in IFT144 (C) causes cranioectodermal dysplasia and is the truncation site in the IFT144$^{\Delta C}$ mutant (Figure 6C). See Table S3.

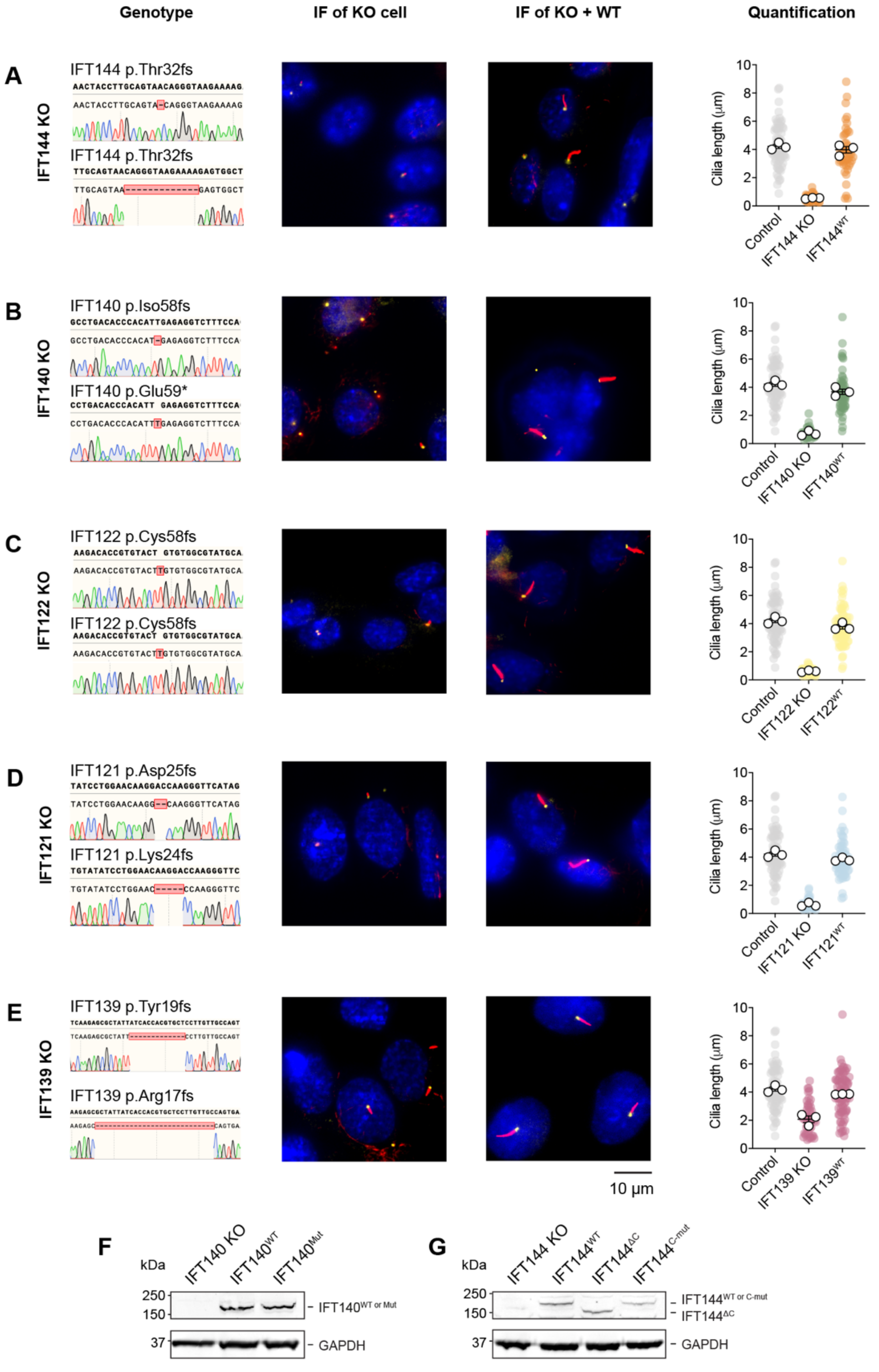


**Figure S4 – CRISPR Knockout of IFT-A Subunits**

(A–E) Left panels, genotype for each IFT-A subunit KO cell line, with indels highlighted by alignment with the reference sequence. Clones were exhaustively sequenced by Sanger sequencing to determine genotype (representative traces shown). IFT144, IFT140, IFT121 and IFT139 KO were compound heterozygous. IFT122 KO was homozygous.

Centre panels, immunofluorescence images of cilia in IFT-A KO cell lines and the corresponding cell lines stably expressing the wild-type (WT) version of the missing subunit. Cells were stained for gamma tubulin (yellow), acetylated tubulin (red) and DAPI (blue). Right panel, quantification of cilia length in control cells (repeated in in each row for comparison), IFT-A KO cell lines, and the corresponding cell lines stably expressing the wild-type version of the missing subunit. Individual data points colored by IFT-A subunit, control in gray. White circles; average for each separate experiment. Lines; mean (± SEM). Control n = 70; KOs IFT144 n = 34, IFT140 n = 51, IFT122 n = 49, IFT121 n = 41, IFT139 n= 50; rescues $IFT144^{WT}$ n= 46, $IFT140^{WT}$ n= 65, $IFT122^{WT}$ n = 74, $IFT121^{WT}$ n = 66, $IFT139^{WT}$ n = 77 cilia analyzed from three separate experiments. One-way ANOVA followed by Kruskal-Wallis test values. Control vs IFT-A KO $p < 0.0001$, IFT-A KO vs $IFT\text{-}A^{WT}$ $p < 0.0001$; control vs $IFT\text{-}A^{WT}$ $p > 0.5$ for all IFT-A proteins.

(F, G) Western blots showing expression of FLAG-tagged IFT140 or IFT144 and associated mutants in IFT140 KO cells (F) or IFT144 KO cells (G). IFT140 and IFT144 detected using anti-FLAG, GAPDH used as loading control.

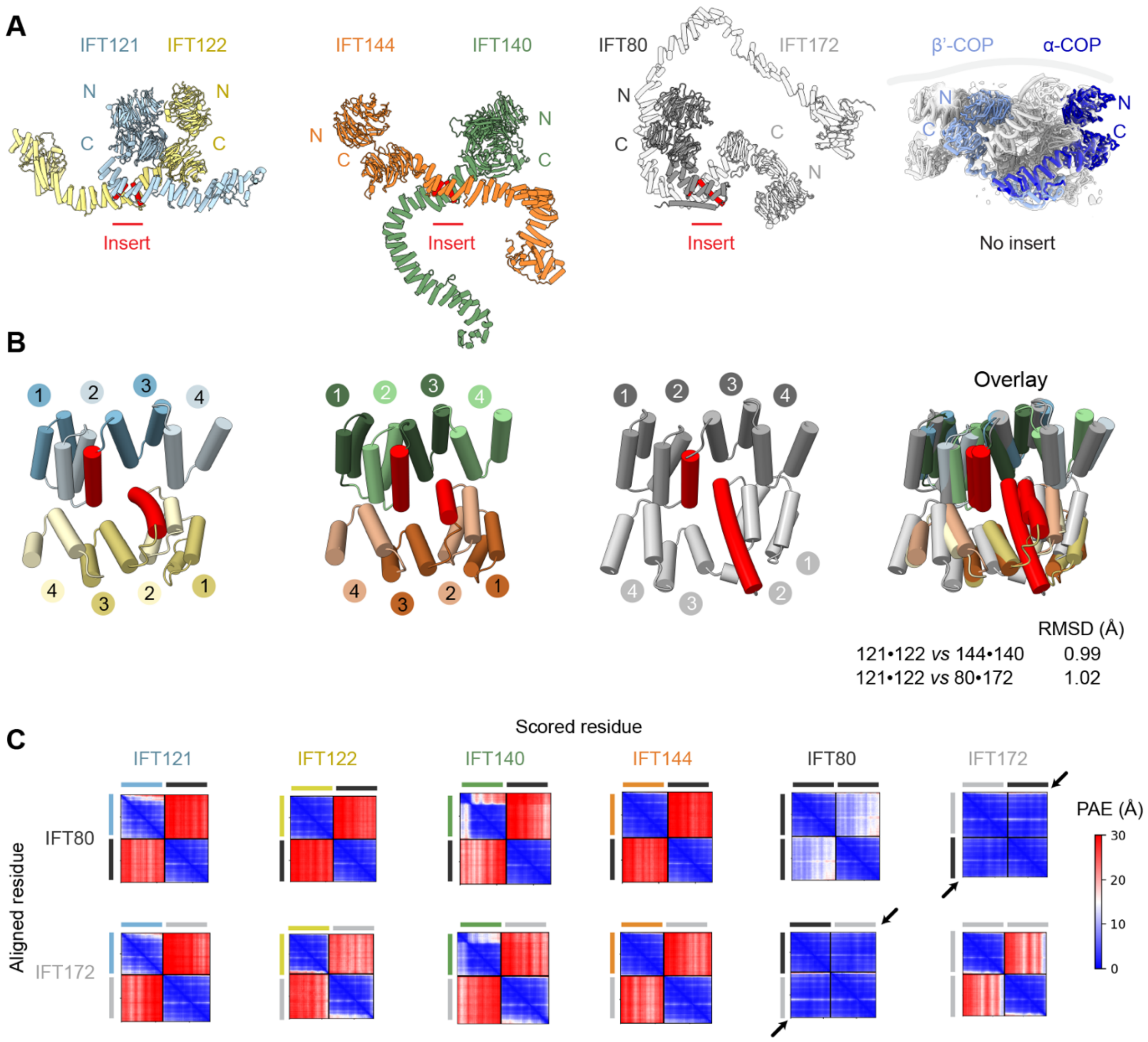


**Figure S5 – Relationship Between IFT-A, IFT-B and Membrane Coat Proteins**

(A) Dimerization interfaces between IFT-A proteins (IFT121•IFT122 and IFT144•IFT140), IFT-B proteins (IFT80•IFT172; AF Multimer prediction) and COPI α•β' (Dodonova et al. 2015). The IFT proteins dimerize via a distinctive α-helix inserted between the second and third TPR of each subunit (red cylinders), whereas COPI α•β' dimerize via a different interface.

(B) Close-up view of TPR interface between IFT proteins. Distinctive α-helix inserted between the second and third TPR of each subunit colored red. TPR repeats numbered. Right, overlay of the structures and root mean square deviation (RMSD) between Cα atom pairs at interfaces (37 pruned atom pairs for 121•122 *vs* 144•140, 57 pruned atom pairs for 121•122 *vs* 80•172).

(C) AF Multimer PAE plots showing specificity of the predicted TPR interface between IFT80 and IFT172. A high confidence (low PAE) interface is predicted between IFT80 and IFT172 (black arrows) but not between IFT80/IFT172 and the other IFT-A proteins (IFT121, IFT122, IFT140 or IFT144). A moderate confidence interface is predicted between IFT80 and IFT80, consistent with the propensity of IFT80 to homodimerize (Taschner et al., 2018).

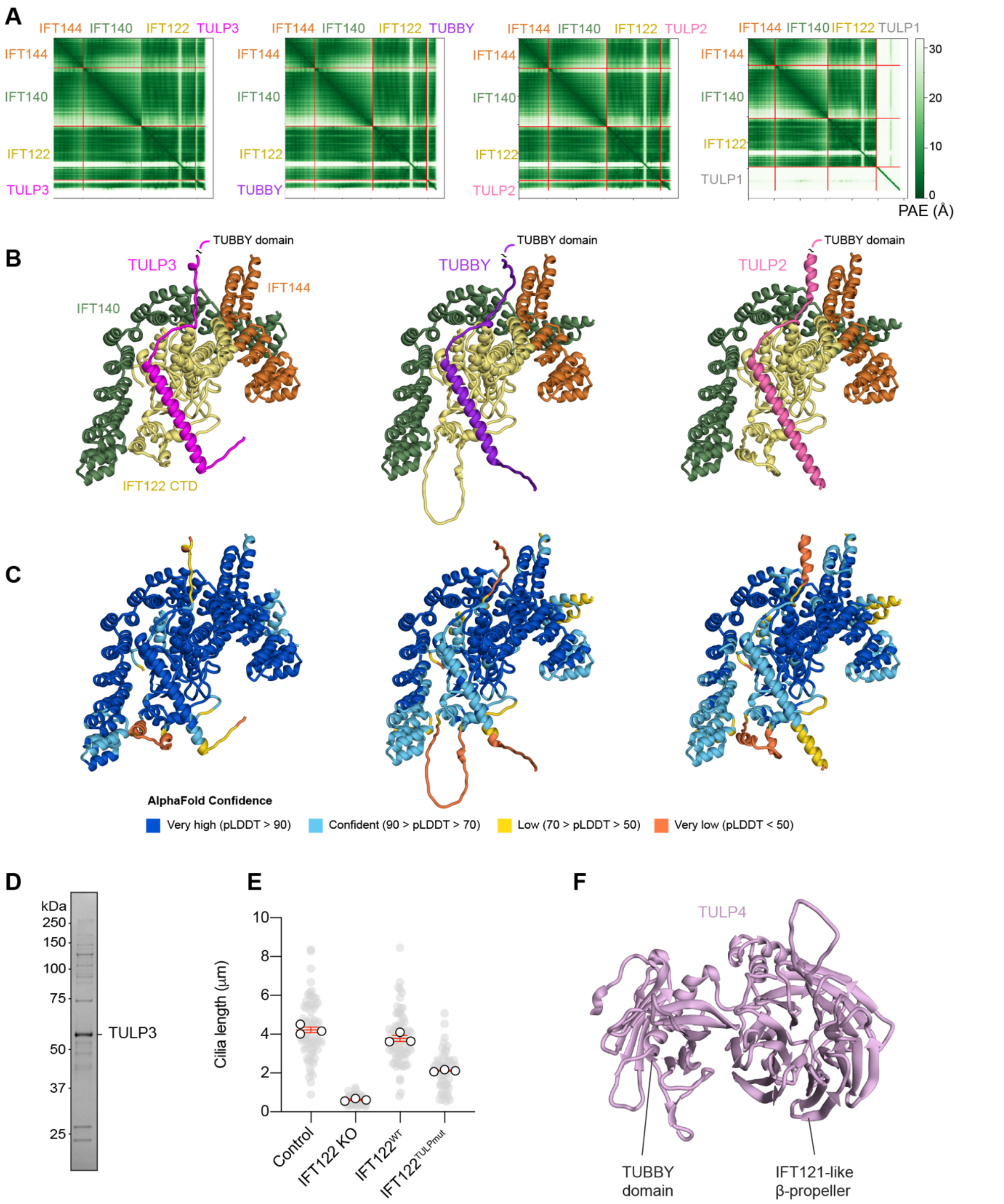


**Figure S6: Analysis of the Interface Between IFT-A and TULP Family Proteins**

(A) AF Multimer prediction of the interface between IFT144•IFT140•IFT122 and TULP3, TUBBY, and TULP2. A non-IFT-A binding TULP protein (TULP1) (Mukhopadhyay and Jackson, 2011) does not a show a predicted interface.

(B) AF Multimer models of the interface between IFT144•IFT140•IFT122 and TULP3, TUBBY, and TULP2, colored by subunit. Note the similar binding mode for each TULP protein.

(C) AF Multimer models of the interface between IFT144•IFT140•IFT122 and TULP3, TUBBY, and TULP2, colored by pLDDT confidence.

(D) SDS-PAGE of purified human TULP3.

(E) Quantification of cilia length in the indicated cells. Short-cilia phenotype in IFT122 KO cells is restored with wild-type IFT122 and partially rescued by IFT122$^{TULPmut}$. Individual data points (gray circles), averages of independent experiments (white circles). Red lines; mean (± SEM). Control n = 70; IFT122 KO n = 49; IFT122 KO + IFT122WT n = 74, + IFT122$^{TULPmut}$ n = 50 cilia analyzed from three separate experiments. One-way ANOVA followed by Kruskal-Wallis test values, control vs IFT122 KO $p < 0.0001$; control vs IFT122WT $p > 0.5$; control vs IFT122$^{TULPmut}$ $p < 0.0001$; IFT122WT vs IFT122$^{TULPmut}$ $p < 0.0001$.

(D) AF model of TULP4; a divergent TULP protein that does not contain an IFT-A binding motif but features a TUBBY domain and IFT121-like β-propeller joined in the same polypeptide, which are predicted to directly interact. Unstructured linker between the domains not shown for clarity.

### Table S1. Cryo-EM data collection

| | IFT-A2 | IFT-A1 |
|---|---|---|
| **Data collection and processing** | | |
| Microscope | Titan Krios (Birkbeck) | Titan Krios (Birkbeck) |
| Detector | K3 | K3 |
| Voltage (keV) | 300 | 300 |
| Nominal magnification | 81,000× | 81,000× |
| Electron exposure ($e^{-}$/Å$^2$) | 48.0 – 49.5 | 48.0 – 49.5 |
| Nominal defocus range (μm) | –1.5 to –3.5 | –1.5 to –3.5 |
| Pixel size (Å) | 1.067 | 1.067 |
| Particles (no.) | 242,645 | 136,617 |
| Global map resolution (Å) | 3.5 | 8 |
| FSC threshold | 0.143 | 0.143 |
| Resolution range (Å) | 3.2–8 | 7–15 |

### Table S2. IFT-A2 atomic model

| | |
|---|---|
| Initial models | - *De novo*<br>- AlphaFold2 Database: IFT139, IFT122, IFT121, IFT43<br>- AlphaFold Multimer: IFT139•IFT121•IFT43 |
| Model composition | |
| Chains | C,D,E,F |
| Nonhydrogen atoms | 20,266 |
| Protein residues | 2,526 |
| R.m.s. deviations | |
| Bond lengths (Å) | 0.012 |
| Bond angles (°) | 1.934 |
| FSC model (0.143 / 0.5) | 3.5 / 4.1 |
| Correlation coefficient (CCmask / box) | 0.67 / 0.77 |
| **Validation** | |
| MolProbity score | 0.90 |
| Clashscore | 0.25 |
| Poor rotamers (%) | 0.09 |
| Ramachandran plot | |
| Favored (%) | 95.65 |
| Allowed (%) | 4.35 |
| Disallowed (%) | 0 |

**Table S3. Disease causing mutations in IFT-A proteins**

| Mutations | Conditions | References |
|---|---|---|
| **IFT144** | | |
| L7P, A30P, V68D, G109E, D159N, L214Ffs5*, F249S, T261Lfs18*, N273D, R272C, G294R, V345G, H481R, D493H, G495R, G495C, L618P, L710S, Q855*, A914D, E1003G, Y1023*, R1103*, R1178Q, E1235K, C1267Y, K1271E | Cranioectodermal dysplasia, short-rib thoracic dysplasia with or without polydactyly, nephronophthisis, Mainzer-Saldino syndrome, Senior-Loken syndrome, retinitis pigmentosa, retinal dystrophy, nonsyndromic asthenoteratospermia | **Bredrup, C. et al.** Am. J. Hum. Genet. 89:634-643 (2011), **Coussa, R. G. et al.** Clin. Genet. 84:150-159 (2013), **Davis, E.E. et al.** Nat. Genet. 43:189-196 (2011), **Fehrenbach, H. et al.** Pediat. Nephrol. 29:1451-1456 (2014), **Halbritter, J. et al.** Hum. Genet. 132:865-884 (2013), Lee, J. et al. Pediat. Nephrol. 30:1451-1458 (2015), **Ni, X. et al.** J. Assist. Reprod. Genet. 37:1431-1439 (2020), **Montolio-Marzo, S. et al.** Europ. J. Med. Genet. 63:104073 (2020), **Zhang, W. et al.** Hum. Mutat. 39:152-166 (2018). |
| **IFT140** | | |
| H24Y, P71L, G140R, L152F, E164*, G212R, I233M, E267G, R280W, I286Kfs5*, V292M, Y311C, C329R, C333Y, A341T, A418P, L440P, W459*, N460Kfs28*, G522E, R576Q, C663W, E664K, R759*, R760*, P726L, E790K, R871C, Q886*, S939P, A974V, G1276E, G1276R, G1305Gfs56*, A1306fs, C1360R , L1399P | Cranioectodermal dysplasia, short-rib thoracic dysplasia with or without polydactyly, nephronophthisis, Mainzer-Saldino syndrome, retinitis pigmentosa, retinal dystrophy | **Bayat, A. et al.** Clin. Dysmorph. 26:247-251 (2017), **Bifari, I. N. et al.** Brit. J. Ophthal. 100:829-833 (2016), **Geoffroy, V. et al.** Hum. Mutat. 39:983-992 (2018), **Hull, S. et al.** Invest. Ophthal. Vis. Sci. 57:1053-1062 (2016), **Khan, A. O., et al.** J. AAPOS 18:203-205 (2014), **Miller, K. A. et al.** PLoS Genet. 9(8): e1003746 (2013), **Perrault, I. et al.** Am. J. Hum. Genet. 90:864-870 (2012), **Schmidts, M. et al.** Hum. Mutat. 34:714-724 (2013), **Xu, M. et al.** Hum. Genet. 134:1069-1078 (2015), **Zhang, W. et al.** Hum. Mutat. 39:152-166 (2018). |
| **IFT139** | | |
| K31fs48*, A44D, F60Y, W150R, K157E, P209L, T231S, A235P, Y255C, A327S, Y347S, R411*, H426D, F440fs443*, P466H, R486Kfs22*, A499T, C518R, C552*, H566R, S591N, D755Y, L795P, Q834*, M844V, R867C, Q869R, D1041N T1103R, Y1167C, M1186V, L1202P, I1208S | Short-rib thoracic dysplasia with or without polydactyly, Joubert syndrome, nephronophthisis, | **Bullich, G. et al.** Nephrol. Dial. Transplant. 32:51-156 (2017), **Cong, E.H. et al.** J Am Soc Nephrol. 25:2435–2443 (2014), **Davis, E.E. et al.** Nat. Genet. 43:189-196 (2011), **McInerney-Leo, A. M. et al.** Clin. Genet. 88:550-557 (2015), **Zhang, H. et al.** Nephrology (Carlton) 4:371-376 (2018), **Zhang, W. et al.** Hum. Mutat. 39:152-166 (2018). |

| | | |
|---|---|---|
| **IFT122** | | |
| W7C, S322F (S263F)[†], Y339Wfs73*, E370Sfs51*, S373F, V391L, V442L, G495R (G436R)[†], V502G (V443G)[†], V553G, G546R, F570C, G572V (G513V)[†], F621C, G623V, L712R, L763P, A1011P (A1062P)[†], Y1077Vfs11* | Cranioectodermal dysplasia, short-rib thoracic dysplasia with or without polydactyly | **Alazami, A.M. et al.** Mol. Genet. Genomic Med. 2:103-106 (2014), **Takahara M., et al.** Hum. Mol. Genet. 27:516-528 (2018), **Tsurusaki, Y. et al.** Clin. Genet. 85:592-594 (2014), **Moosa, S. et al.** J. Med. Genet. A 170A:1295-1301 (2016), **Silveira, K. C. et al.** Am. J. Med. Genet. A 173:1186-1189 (2017), **Walczak-Sztulpa, J. et al.** Am. J. Med. Genet. 173A:1364-1368 (2017). |
| **IFT121** | | |
| I9Tfs7*, W261R, Q276*, W311L, R478K, G501Kfs1*, L520P, Q527*, R545*, E626G, L641*, D841V, A875T, P964Lfs15*, H1031Y, Y1068C, W1153C | Cranioectodermal dysplasia, short-rib thoracic dysplasia with or without polydactyly | **Bacino, C. A. et al.** Am. J. Med. Genet. 158A:2917-2924 (2012), **Duran, I. et al.** Cilia 6:7 (2017), **Gilissen, C. et al.** Am. J. Hum. Genet. 87:418-423 (2010), **Lin, A. E. et al.** Am. J. Med. Genet. 161A:2762-2776 (2013), **Mill, P. et al.** Am. J. Hum. Genet. 88:508-515 (2011), **Smith, C. et al.** Am. J. Med. Genet. 170A:760-765 (2016), **Toriyama, M. et al.** Nature Genet. 48:648-656 (2016), **Walczak-Sztulpa, J. et al.** Am. J. Med. Genet. 173A:1364-1368 (2017), **Zhang, W. et al.** Hum. Mutat. 39:152-166 (2018). |
| **IFT43** | | |
| M1K, E34K, W174R (W179R)[†] | Short-rib thoracic dysplasia with or without polydactyly | **Biswas, P. et al.** Hum. Mol. Genet. 26:4741-4751 (2017), **Duran, I. et al.** Cilia 6:7 (2017). |

[†] Non-canonical isoform numbering in brackets.

### Supplemental Movie Legends

**Movie S1 – Multi-body analysis of IFT-A.**
Conformational flexibility of IFT-A1 module with respect to IFT-A2 (fixed frame of reference). Three principal components of motion, representing rotations about three orthogonal axes (determined using RELION multi-body analysis; Nakane et al., 2018), are shown sequentially.

**Movie S2 – Structure of the IFT-A Complex.**
Depiction of the IFT-A complex and pseudo-atomic model of the IFT-A polymer.
